# Proteome and phospholipidome interrelationship of synovial fluid-derived extracellular vesicles in equine osteoarthritis: An exploratory ‘multi-omics’ study to identify composite biomarkers

**DOI:** 10.1101/2023.08.02.551609

**Authors:** Emily J Clarke, Laura Varela, Rosalind E Jenkins, Estefanía Lozano−Andrés, Anna Cywińska, Maciej Przewozny, P. René van Weeren, Chris H.A. van de Lest, Mandy Peffers, Marca H.M. Wauben

**Author notes:** Corresponding authors: *E-mail address:* (M.H.M.Wauben), *E-mail address:* (M. Peffers). E.C. and L.V. contributed equally to this work. M.P. and M.H.M.W. contributed equally to this work.

## Abstract

Osteoarthritis causes progressive joint deterioration, severe morbidity, and reduced mobility in both humans and horses. Currently, osteoarthritis is diagnosed at late stages through clinical examination and radiographic imaging, hence it is challenging to address and provide timely therapeutic interventions to slow disease progression or ameliorate symptoms. Extracellular vesicles are cell–derived vesicles that play a key role in cell–to–cell communication and are potential sources for specific composite biomarker panel discovery. We here used a multi–omics strategy combining proteomics and phospholipidomics in an integral approach to identify composite biomarkers associated to purified extracellular vesicles from synovial fluid of healthy, mildly and severely osteoarthritic equine joints. Although the number of extracellular vesicles was unaffected by osteoarthritis, proteome profiling of extracellular vesicles by mass spectrometry identified 40 differentially expressed proteins (non–adjusted p<0.05) in osteoarthritic joints associated with 7 significant canonical pathways in osteoarthritis. Moreover, pathway analysis unveiled changes in disease and molecular functions during osteoarthritis development. Phospholipidome profiling by mass spectrometry showed a relative increase in sphingomyelin and a decrease in phosphatidylcholine, phosphatidylinositol, and phosphatidylserine in extracellular vesicles derived from osteoarthritic joints compared to healthy joints. Unsupervised data integration revealed positive correlations between the proteome and the phospholipidome. Comprehensive analysis showed that some phospholipids and their related proteins increased as the severity of osteoarthritis progressed, while others decreased or remained stable. Altogether our data show interrelationships between synovial fluid extracellular vesicle–associated phospholipids and proteins responding to osteoarthritis pathology and which could be explored as potential composite diagnostic biomarkers of disease.

## 1. Introduction

Osteoarthritis (OA) is the most prevalent arthritic phenotype and is one of the most important causes of perception of pain and loss of quality of life in the older population [1]. OA has often been classified as a chronic degenerative joint disease resulting from a process of wear and tear. However, OA has an important inflammatory component, the mediators of which trigger an aberrant remodelling of joint structures inside the afflicted articulation [2]. These may include synovial membrane dysfunction, abnormal bone proliferation, and subchondral bone sclerosis [3]. Age, gender, obesity, genetics, inactivity, joint loading, aberrant morphology and alignment, previous injuries, and muscle weakness are the most prevalent risk factors for OA [4]. OA in horses is a major cause of lameness, with over 60% of lameness cases associated with a clinical diagnosis of OA [5]. This results in impaired mobility, pain, poor performance, and early retirement, making equine OA a serious welfare issue that also leads to significant economic losses for the equine industry [6].

Previously, it has been shown that human and equine osteoarthritic pathogenesis follows a similar route from initial injury to disease progression and outcome, and as such, the horse is widely regarded as a clinically relevant model for musculoskeletal disease in humans [7]. In addition, the horse’s articular cartilage biology is anatomically comparable to that of humans with respect to both composition and thickness [8]. The horse as a model for disease offers numerous further benefits, including the applicability of advanced diagnostic methodologies, such as magnetic resonance imaging (MRI) and arthroscopy, as well as serial sampling of biological material for analysis making it possible to monitor disease development, disease progression and response to intervention in great detail [7].

Presently, OA pathophysiology is not fully understood. The diagnosis is commonly based on clinical examination and radiographic imaging and, due to the insidious character of the disorder is often made at late stages when cartilage damage is already substantial and far exceeds the tissue’s capacity for intrinsic repair [9]. Therefore, it is paramount to identify biomarkers of disease that can be used to develop diagnostic tests that are both sensitive and specific for early OA, which could ultimately enable a timelier management of therapeutic interventions and decelerate disease progression.

In recent years, the concept of composite biomarkers has become popular; by definition, they are a non–linear combination of multiple measurements used to diagnose disease or predict outcomes [10]. Thus far, they have been used in neurological diseases such as Alzheimer’s disease and bipolar disorder [11], often using neuronal networks, artificial intelligence or machine learning algorithms. As such, extracellular vesicles (EVs) can be considered a biological source for composite biomarker discovery.

EVs are nanoscale–sized vesicles with a phospholipid bilayer membrane secreted by cells and specialised in restoring homeostasis or facilitating intercellular communication [12]. Furthermore, EVs transport bioactive molecules that can elicit a response in recipient cells, resulting in physiological and phenotypic changes [13; 14]. They are present in tissues and body fluids, such as blood, urine and synovial fluid (SF) [15–17]. It has been proposed that EVs may play a vital role in cartilage homeostasis and in the propagation of OA by promoting inflammation and regulating extracellular matrix (ECM) turnover [18–21]. EVs are found in abundance in SF due to its close proximity to EV–secreting sources, such as native cells found within the joint space and periarticular tissues, including but not limited to chondrocytes and synoviocytes [22]. For joint disorders such as OA, SF is thus the most appropriate source of biochemical information [20; 21; 23].

The translation of EV biomarkers to the clinic has been pioneered in the fields of cancer and neurodegenerative diseases [24; 25]. Nowadays, EVs are increasingly seen as a source for biomarker discovery for various disorders, including joint disease [18; 20; 26]. A comprehensive understanding of the molecular composition of EVs and their role in disease requires the interpretation of molecular intricacy by accounting for multiple biological levels, such as the proteome and phospholipidome [27; 28]. Such a comprehensive experimental and data analysis approach provides a more thorough understanding of the complete spectrum of molecular changes contributing to cellular response, disease development and pathogenesis and is helpful for the identification of naturally occurring composite biomarkers. Recent studies in ovarian cancer [29] and Alzheimer’s disease [30] have identified a relationship between the proteome and phospholipidome of EVs.

The present exploratory study exploited omics–based technologies to analyse the proteome and phospholipidome of SF–derived EVs (SF–EVs) to 1) enable comprehensive profiling of a healthy state versus clinically diagnosed mild and severe OA in horses and 2) identify candidate composite diagnostic biomarkers of OA.

## 2. Materials and Methods

### 2.1 Ethical Considerations

Equine SF was collected from horses presenting at the EQI VET SERWIS clinic in Buk, Poland, with various disorders of the locomotor system before the intra–articular application of a local analgesic as a standard part of the clinical lameness examination. Sample collection was approved by the University of Liverpool’s Veterinary Research Ethics Committee (VREC1180). Ethical approval was not required in Poland, as the procedures were considered non–experimental clinical veterinary practices, in accordance with Polish and EU law (Dz. U. 2015 poz. 266 and 2010–63–EU directive).

### 2.2. Sample collection

SF was collected via aseptic arthrocentesis from the metacarpophalangeal joint into a plain Eppendorf tube. Samples were spun at 2,540 ×g at 4°C for 5 minutes. The supernatant was then transferred to a new Eppendorf tube, snap–frozen in liquid nitrogen and stored at –80°C.

OA was diagnosed based on clinical examination, including subjective lameness scoring according to the American Association of Equine Practitioners (AAEP) lameness scale, radiographic imaging and diagnostic analgesia. Classification as “mild OA” – a mild form of the disease (minor lesions, e.g. small osteophytes and limited subchondral bone sclerosis) and “severe OA” – a severe form of the disease (bone deformation, clear subchondral bone sclerosis, narrowing of joint space and formation of larger osteophytes) was based on radiography; radiographic examples of each OA phenotype are shown in Suppl. Table 1. The horses with no lesions in the joints (which featured locomotor abnormalities caused by disorders of other, unrelated structures) were classified as horses with healthy joints. Three biological replicates were pooled per sample resulting in 5mL of SF. Pooled samples came from horses with healthy joints or with the same disease severity. Donors for the pooled samples were randomised with respect to age and sex. A total of 42 donors were used, resulting in 14 pooled samples (Healthy joints n=7, mild OA n= 4, and severe OA n= 3).

### 2.3. Extracellular vesicle isolation

#### 2.3.1. Differential centrifugation

EVs from SF were isolated using a published and validated method [16]. First, the pooled cell–free SF samples (5 mL) were incubated at 37 °C for 15 minutes with HYase (5 mg/mL; HYase type II from sheep testes, Sigma– Aldrich, St. Louis, USA) while vortexing every 5 minutes. Next, (protein) aggregates and debris were removed by centrifuging at 1,000 ×g for 10 minutes at RT (Avanti J–15R; Beckman Coulter Inc., Brea, USA). Next, the supernatants were transferred into SW40 tubes (Beckman Coulter Inc., Brea, USA) and mixed with phosphate–buffered saline (PBS) to a volume of 12 mL, and centrifuged at 10,000 ×g for 35 minutes (8,900 RPM; RCF average 10,003 ×g; RCF max 14,088 ×g; *κ*–Factor 2,771), followed by 100,000 ×g for 65 minutes (28,000 RPM; RCF average 99,004 ×g; RCF max 139,439 ×g; *κ*–Factor 280). The 40 Ti Beckman Coulter rotors were used in an Optima™ L–90K or Optima™ XPN–80 ultracentrifuges. EV pellets were resuspended in 300 *μ*L PBS+0.1% Bovine Serum Albumin (BSA) EV-depleted (previously depleted of EVs and protein aggregates).

#### 2.3.2. Sucrose density gradient

EV pellets were thoroughly mixed with 1.2 mL 2.5 M sucrose solution (J.T. Baker; Phillipsburg, USA) in a new SW40 tube and overlaid with fourteen sucrose solutions of decreasing density (from 2 M to 0.4 M), creating a discontinuous sucrose gradient. Gradients were centrifuged at 200,000 ×g for 16 hours at 4 °C in an SW40–Ti rotor (39,000 RPM; RCF average 192,072 ×g; RCF max 270,519 ×g; *κ*–factor 144.5). Twelve fractions of 1 mL were collected from the top (lowest density) to the bottom (highest density). Fraction densities were determined by refractometry. EV–containing fractions (validated in [16; 21]) were pooled (densities between 1.10–1.16 g/mL) and pipetted into SW32 ultracentrifuge tubes for lipidomics and SW60 for proteomics analysis. EVs were pelleted by centrifugation for 95 minutes at 120,000 ×g at 4°C (SW32 Ti. 32,000 RPM; RCF average 127,755 ×g; RCF max 174,899 ×g; *κ*–Factor 204, or SW60 Ti, 35,000 RPM; RCF average 125,812 ×g; RCF max 165,052 ×g; *κ*–Factor 133). Subsequently, Evs for lipidomics were resuspended in 100 *μ*L of PBS. For proteomics, dried EV pellets were snap–frozen immediately at – 20°C for later analysis.

Relevant data regarding the experimental details for EV isolation and characterisation have been submitted to the EV–TRACK knowledgebase (EV–TRACK ID: EV230607) [31].

### 2.4. Single–EV–based high–resolution flow cytometry

#### 2.4.1. Labelling of EV pellets with PKH67

Generic fluorescent staining of EVs was performed with the PKH67 labelling kit (Sigma–Aldrich, St. Louis, USA) as previously described [21; 32], with minor modifications indicated below. EV pellets were resuspended in 20 *μ*L PBS+0.1% BSA EV–depleted with 30 *μ*L diluent C. Then, 50 *μ*L of diluent C with 1.5 *μ*L of PKH67 dye were added. The staining process was halted by adding 50 *μ*L of EV–depleted RPMI/10% FBS (Roswell Park Memorial Institute/fetal bovine serum) after 3 min of incubation at room temperature. Next, the labelled EVs were combined with 2.5 M sucrose to continue the previously described density gradient ultracentrifugation process for EV separation in section 2.3.2. Throughout the whole PKH67 labelling and sucrose gradient ultracentrifugation process, a procedural control sample (20 *μ*L PBS+0.1% BSA EV–depleted +30 *μ*L diluent C, without sample EVs) was used as a control sample for high–resolution FCM.

#### 2.4.2. Single EV–based high–resolution FCM analysis

An optimised BD Influx jet–in–air– flow cytometer (Becton Dickinson Biosciences, San Jose, CA, USA) fully tailored for single EV analysis was employed [32; 33]. In short, a workspace was loaded that had the optimised PMT parameters and pre–defined gates for the detection of 200 nm yellow–green (505/515) FluoSphere beads (Invitrogen, F8848). After the fluid stream and lasers were aligned, the 200 nm bead population had to fulfil the requirements of pre–defined mean fluorescent intensity and scatter values inside these gates while exhibiting the lowest coefficient of variation for side scatter, forward scatter, and FL–1 fluorescence. All measurements in this study used the same fluorescent threshold level, which was established by running a clean PBS sample and allowing an event rate of < 20 events/ second. All samples were run for a fixed period of 30 seconds. The EV concentration was calculated based on the number of fluorescent events detected and normalised for the flow rate of 12.8 *μ*L/minute, dilution factor and measurement time. The final EV concentration per mL of SF was determined for the EV–enriched sucrose fractions F7–F10 (densities 1.10 g/mL–1.16 g/mL), adjusted based on the SF starting volume. The procedural control sample revealed no noteworthy background events (<500 events/ 30 secs) in gradient fractions of interest (data not shown).

The BD FACS Software v1.0.1.654 (BD Biosciences, San Jose, CA, USA) was used to collect the data, and the FlowJo v10.07 software (FlowJo, Ashland, OR, USA) was used for analysis. The MIFlowCyt author checklist can be found on Suppl. Table 2 and the MIFlowCyt–EV framework on Suppl. Table 3 [34].

### 2.5. Lipidomic analysis

#### 2.5.1. Lipid extraction

Lipids were extracted following the Bligh & Dyer method [35] with slight modifications. First, 0.7 mL of fresh deionised water was mixed with 100 *μ*L of samples, 2 mL methanol, and 1 mL chloroform. The samples were incubated for 20 minutes, then 2 mL chloroform and 2 mL of deionised water were added, and the mixture was vortexed. The resultant hydrophilic and hydrophobic phases were separated by centrifugation at RT for 5 minutes at 2,000 ×g. The hydrophobic bottom phase was transferred to a new conical glass tube. The extraction of the remaining sample (hydrophilic phase) was repeated with an additional 2 mL of chloroform to ensure that all lipids were collected. The samples were dried under nitrogen gas injection and stored in a nitrogen atmosphere at –20 °C. During the lipid extraction, one sample from healthy joints was lost. Therefore, n=6 SF–EV samples of the group with healthy joints were used for all lipidomics analyses and subsequent omics integration.

#### 2.5.2. Mass spectrometry lipidomics

Dried lipid pellets were resuspended in 30 *μ*L chloroform/methanol (1:1) and analysed as described previously [21; 36]. A quality control sample composed of all samples in the same ratio together with the SPLASH® Lipidomix® Mass Spec Standard (Avanti Polar Lipids, Inc., Alabaster USA) was created for subsequent lipid quantification and added in the MS run. A hydrophilic interaction liquid chromatography (HILIC) column (2.6 *μ*m Kinetex HILIC 100, 50 × 4.6 mm, Phenomenex, Torrance, USA) was loaded with 10 *μ*L of lipid extract. Gradient elution on an Agilent 1290 InfinityII UPLC (Agilent, CA) separated the lipid classes. Solvent A consisted of acetonitrile/acetone (9:1) with 0.1% formic acid, and solvent B was composed of acetonitrile/H2O (7:3) with 0.1% formic acid and 10 mM ammonium formate. The gradient profile was: minute 0 to 1: 100% A; minute 1 to 3: 50% A + 50% B; minute 3 to 5: 100% B, with a 1 mL/min flow rate. Without further re–equilibration of the column, samples were injected.

The samples were analysed with a Fusion Orbitrap MS (ThermoFisher Scientific, Waltham, USA) via a heated electrospray ionisation (HESI) source with the following parameters: negative ion spray voltage, 3.6 kV; aux sheath gas flow rate, 54Arb; gas flow rate, 7Arb; sweep gas flow rate, 1Arb; ion transfer tube temperature, 350°C; vaporiser temperature, 450°C; scan range, 350–950 m/z at a resolution of 120,000.

#### 2.5.3. Lipid annotation

The msconvert ProteoWizard [37] was used to convert the RAW format to mzML with the “peakPicking filter vendor msLevel = 1–” parameter selected. The package XCMS version 3.10.2 [38] was used to perform LC/MS peak–picking, sample–grouping, and retention time correction on the mzML files running under R version 4.1.2. The identified LC/MS peaks (features) were annotated based on retention time, exact m/z–ratio, and, if present in at least 3 out of 13 pooled samples (Healthy n=6, OA n=4, Advanced OA n=3), MS peaks were annotated using an in–silico phospholipid database. The features were also selected to account for isotope distribution and adducts. The RAW and mzML converted mass spectrometry data is deposited in the YODA repository of Utrecht University [39].

### 2.6. Proteomic analysis

#### 2.6.1. Protein extraction

EV pellets were resuspended in 200 *μ*L of urea lysis buffer (6 M Urea; Sigma–Aldrich, Dorset, United Kingdom), 1 M ammonium bicarbonate (Fluka Chemicals Ltd., Gillingham, UK) and 0.5% sodium deoxycholate (Sigma–Aldrich, Dorset, United Kingdom). Samples were sonicated at 5 *μ*m for 3 × 10 secs per sample, with 1–min rest on ice between each sonication round as previously described [23].

#### 2.6.2. SDS PAGE & silver stain

Sodium dodecyl sulfate–polyacrylamide gel electrophoresis (SDS–PAGE) was used to separate proteins from EV protein extracts. 7.5 *μ*L of 2x Novex™ Tris–Glycine SDS Sample Buffer (ThermoFisher Scientific, Paisley, UK), supplemented with 8% of 2–Mercaptoethanol (Sigma–Aldrich, Dorset, UK), was added to 7.5 *μ*L of sample SF–EV protein lysate. Samples were mixed and heated at 100°C for 10 min to denature proteins, then placed on ice. Electrophoresis was performed using A NuPAGE™ 4 to 12%, Bis–Tris gel (ThermoFisher Scientific, Paisley, UK) and 1x Nu-PAGE® MES Running Buffer (ThermoFisher Scientific, Paisley, UK) (diluted from the 20x stock in ultrapure water). Samples were loaded onto the gel alongside the Novex™ Sharp Pre–stained Protein Standard ladder (ThermoFisher Scientific, Paisley, UK). Gels were run at 100V until completion of electrophoresis and visualised using silver stain (Thermofisher Scientific, Paisley, UK) according to the manufacturer’s guidelines as previously done [23].

#### 2.6.3. On bead digestion

Hydrophilic and hydrophobic magnetic beads were used for EV protein digestion in order to remove the urea lysis buffer that was not mass spectrometry compatible. Beads were suspended within the lysed EV samples in order for extracted EV proteins to bind to the surface of the bead, and thus a tryptic digest was performed on bead.

Specifically, 95 *μ*L of lysed and sonicated equine SF–EV were treated with 5 mM dithiothreitol (DTT) (Sigma–Aldrich, Dorset, UK) 100 mM at 60°C and shaken at 1000 rpm for 30 mins on an orbital shaker. Iodoacetamide (Sigma–Aldrich, Dorset, UK) was then added to a final concentration of 20 mM, and the samples were incubated at room temperature in the dark for 30 min. Following this, 5 mM DTT was added to each sample and incubated at room temperature for 15 min. Hydrophilic and hydrophobic magnetic carboxylate Speed-Beads (SP3 beads, total of 12 *μ*L) (Cytiva, Massachusetts, US) were added to each sample, followed by 120 *μ*L ethanol (Sigma– Aldrich, Dorset, UK). Samples were then incubated at 24°C and shaken at 1000 rpm for 1 h. The beads were separated from samples using a magnetic stand, washed thrice with 180 *μ*L 80% ethanol and resuspended in 100 mM ammonium bicarbonate (Fluka Chemicals Ltd., Gillingham, UK, 4 *μ*g). Trypsin/LysC (2.4 *μ*g) (Promega) was added to each sample. Samples were placed in a sonicator bath and sonicated for 30 secs to disaggregate the beads before being incubated overnight at 37°C and shaken at 1000 rpm. Beads were removed from the samples using the magnetic stand, and the supernatants were acidified by adding 1 *μ*L 10% trifluoroacetic acid (Sigma– Aldrich, Dorset, UK). Samples were then desalted using an Agilent mRP–C18 column, dried in a SpeedVac and resuspended in 0.1% formic acid. The UV absorbance measured during desalting was used to normalise the loading for mass spectrometry analysis, with a final volume of 5 *μ*L being loaded on the nano–LC column as previously described [23].

#### 2.6.4. Data–dependent acquisition for the generation of an equine SF EV spectral library

Equine SF was pooled using samples from the metacar-pophalangeal joint from our equine musculoskeletal biobank (VREC561) and samples collected in previous studies from the carpal and metacarpal joint of healthy horses as well as those with OA, resulting in a total of 11 ml SF. These samples were analysed as previously described in order to generate the necessary reference library [40].

#### 2.6.5. Data–independent acquisition proteomics (SWATH)

A data–independent proteomic approach was utilised in the form of Sequential Windowed Acquisition of all theoretical fragments (SWATH). Data were acquired using the same 2 h gradient as the library fractions [23]. SWATH acquisitions were performed on a Triple TOF 6600 (Sciex) via an Eksigent nanoLC 415 fitted with an ACQUITY UPLC Peptide BEH C18 nanoACQUITY Column (Waters, UK) and a bioZEN 2.6 *μ*m Peptide XB–C18 (FS) nano column (250 mm x 75 *μ*m, Phenomenex). Data were acquired using 100 windows of variable effective isolation width to cover a precursor m/z range of 400–1500 and a product ion m/z range of 100–1650. Scan times were 50 ms for TOF–MS and 36 ms for each SWATH window, giving a total cycle time of 3.7 secs. Retention time alignment and peptide/protein quantification were performed by Data–Independent Acquisition by Neural Networks (DIA–NN), using the same reference horse proteome as described in section 2.6.3 to reannotate the library. A precursor FDR of 1%, with match between runs and unrelated runs was selected. The mass spectrometry proteomics data were deposited to the ProteomeXchange Consortium via PRIDE proteome exchange [23] (identifier PXD042765). Both proteomics and lipidomics datasets have been submitted to vesiclepedia [41].

### 2.7. Statistical analysis

#### 2.7.1. Proteomics

Statistical analysis of proteomics data was carried out using the R statistical programming environment or Metaboanalyst [42]. The data were quality controlled; proteins with complete observations were normalised using probabilistic quotient normalisation (PQN) and log–transformed (base 10) for downstream analysis. Unsupervised multivariate analysis in the form of principal component analysis (PCA) was performed, along with heat map analysis using analysis of variance (ANOVA) and Pearson distance. Statistical significance by ANOVA was attributed to proteins with a p–value <0.05. Following ANOVA, a fold change analysis was conducted.

#### 2.7.2. Lipidomics and omics data integration

For lipidomics analysis, the data were normalised based on the sum of total lipids per pool sample – i.e. each lipid value in a pooled sample was divided by the total sum of lipids in the same pool sample and multiplied by 0.01; thus, the relative abundances sum up to 100. A minimum of three biological–pool replicates were used for statistical analyses.

Data analysis was run with R version 4.1.2 [43]. Pareto scaling was performed for the PCA, thus dividing each variable by the square root of its standard deviation. Heatmap and cluster analysis was performed on Spearman correlations with a set speed of two – among the 50 most abundant lipid species in all sample groups – using the R–package ComplexHeatmap v1.12.0 [44].

Data integration was performed with the R package mixOmics v6.12.2. [45] on lipidomic and proteomics data normalised by the sum (as described for lipidomics analysis) followed by R scaling and centring, which determines the vector’s mean and standard deviation, deducts the mean from the vector and divides it by the standard deviation. An unsupervised sparse Partial Least Squares (a linear, multivariate regression method for data reduction to assess the relationship between independent and dependent variables) was used to integrate the datasets. The relevance network plot was set with a correlation cut off of 0.7 to allow readability of the displayed proteins and phospholipids. Differences between the proposed proteins and phospholipid percentages for the composite biomarker were analysed with the rank–based non–parametric Kruskal–Wallis test, followed by the multiple pairwise comparisons with Dunn’s test. Significance was defined as p–value < 0.05. Statistical tests were done with GraphPad Prism 9.

### 2.8. Functional enrichment analysis

Functional enrichment analysis was performed on proteomic data using Ingenuity Pathway Analysis (IPA; Qiagen, Hilden, The Netherlands) in order to provide functional analyses, networks, canonical pathways, and related molecular and pathological functions by using protein p–values, including those differentially expressed (values of less than 0.05), and associated log2 fold change. UniProt_Horse accession codes were used as protein identifiers, and the Qiagen Ingenuity Knowledge Base was used as a reference for exploratory pathway analysis. For network generation, default settings were used to identify molecules whose expression was significantly differentially regulated. These molecules were overlaid onto a global molecular network contained in the Ingenuity Knowledge Base. Networks of ‘network–eligible molecules’ were then algorithmically generated based on their connectivity. The functional analysis identified the biological functions and diseases that were most significant to the data set. A right-tailed Fisher’s exact test was used to calculate p–values. Canonical pathway analysis identified the pathways from the IPA library that were most significant to the data set. Analysis was performed on all proteomics data, comparing healthy, mild OA and severe OA groups, and those proteins correlated to phospholipids.

## 3. Results

### 3.1. EV characterisation

#### 3.1.1. Synovial fluid–derived extracellular vesicle numbers do not significantly differ between osteoarthritic and healthy phenotypes

Recently we found that an inflammatory insult in the joint, such as LPS, can strongly affect the quantity of SF–EVs [21]. Therefore, we investigated, using the same technology, if the quantity of SF–EVs was altered as a result of OA using samples from equine patients with radiographically diagnosed OA and comparing these with samples from healthy joints. The quantity of EVs was assessed by single–EV fluorescence–based flow cytometric analysis of PKH–labelled EVs [21; 33] on 3 representative samples of the group with healthy joints, 2 samples of the mild OA group and 1 of the severe OA group. The PKH+ events were measured in individual sucrose fractions ranging from 1.08 to 1.18 g/mL. The peak of fluorescent events was identified in the densities from 1.10 to 1.16 g/mL (Fig. 1A); those were considered the EV–enriched fractions and were used for calculating EV numbers (Fig. 1B). We did not observe statistically significant differences between the numbers of SF–EV from the healthy joints (3.1×10^8^ per mL SF ± 6.6×10^7^; mean ± SD) and OA samples (4.0×10^8^ per mL ± 9.4×10^7^; mean ± SD).

**Figure 1:**
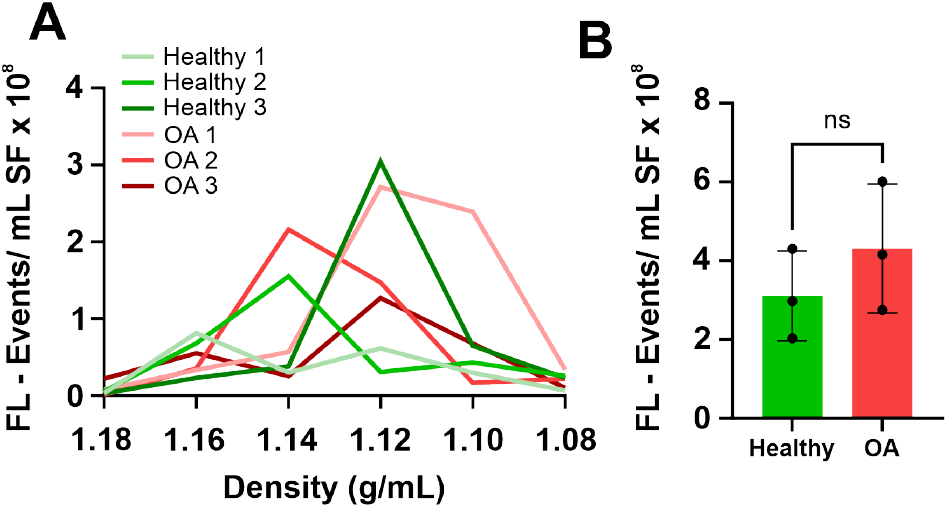
Quantitative flow cytometric analysis of EVs isolated from equine joints with a healthy or osteoarthritic phenotype. **A)** Single EV–based high–resolution FCM of representative healthy SF–EVs (n=3) and OA SF–EVs (n=2) from the mild OA group and n=1 from the severe OA group. Sucrose density gradient fractions containing EVs labelled with the lipophilic dye PKH67 were measured for 30 seconds. The majority of EVs floated at densities of 1.16–1.10 g/mL. FL – Events: Fluorescent Events. **B)** EV concentration in SF was calculated as the sum of single fluorescent events measurements (PKH67+ events) in EV–containing sucrose gradient densities (1.16 to 1.10 g/mL). Mean ± SD. ns: non–significance by Student’s t–test. The uppermost point in the OA group reflects the severe OA phenotype.

### 3.2. Lipidomic Analysis

#### 3.2.1. Synovial fluid–derived extracellular vesicle phospholipid profiles change during the development of osteoarthritis

Previously we had observed a drastic change in the phospholipidome following an inflammatory stimulus [21]; here we analysed whether the phospholipid profile of the SF–EVs was modified as a result of OA. The phospholipidome profile of the SF–EV from healthy joints, mild OA, and severe OA equine patients was determined through a bioinformatics analysis that uncovered 280 lipid species after lipid annotation (and background adjustment), isotope and adduct correction and normalisation by the cumulative sum to unity (Supplementary Figure 1). A PCA, an unsupervised dimensionality reduction method, revealed a combined explained variance of 69% with the first and second principal components (Fig. 2A).

**Figure 2:**
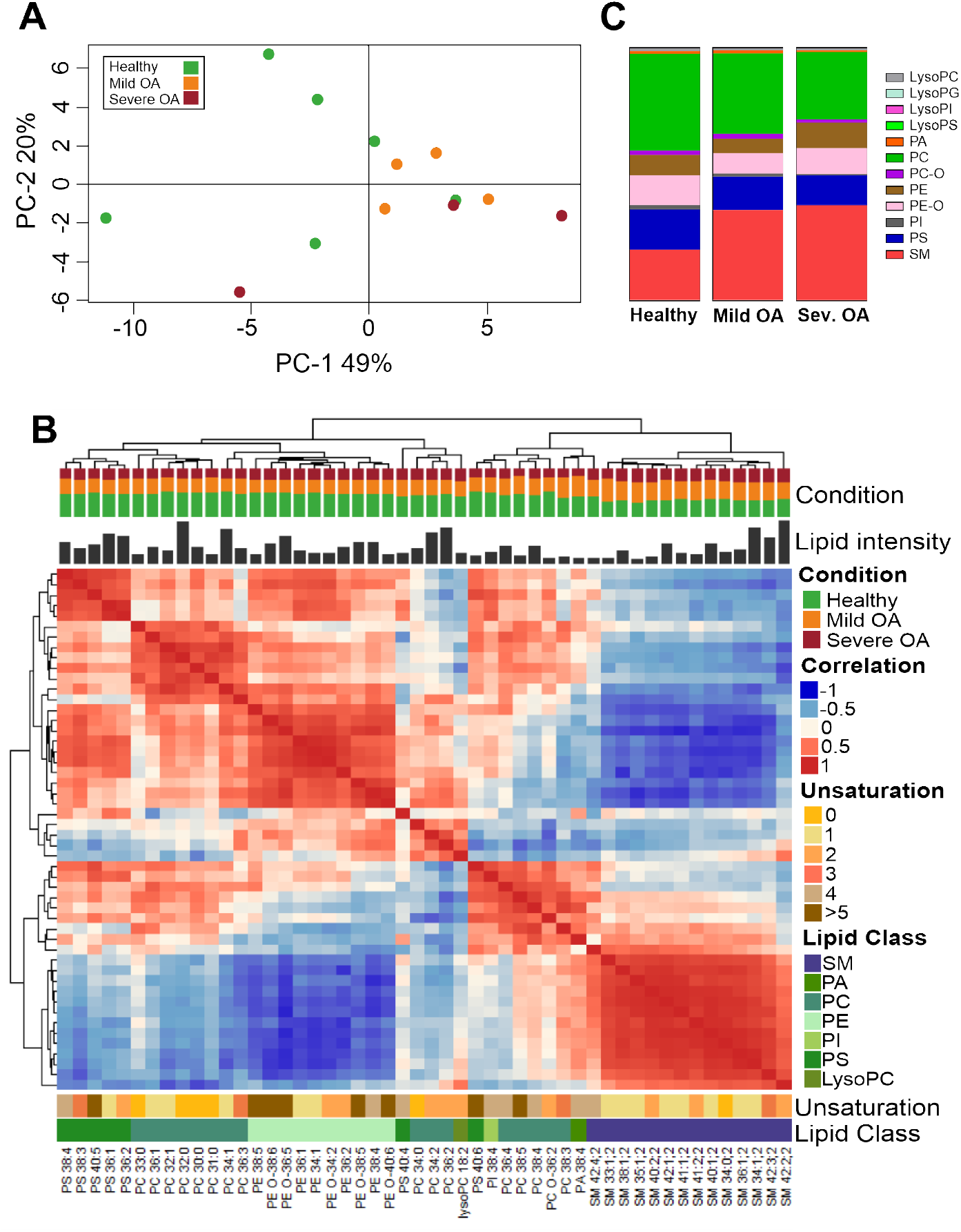
Lipidomic profile of equine synovial fluid–derived EVs from healthy joints or from mild OA or severe OA patients. Healthy samples (n=6), mild OA (n=4), severe OA (n=3). Each sample is comprised of a pool of three different animals. Lipids were extracted from EVs isolated by differential centrifugation up to 100,000g, followed by purification with sucrose density gradients. **A)** Principal component analysis of lipids isolated from the three different clinical groups. The principal components (PC)–1 and –2 explain 49% and 20% of the variance, respectively. Healthy samples (green circle), mild OA (orange circle), and severe OA (red circle). **B)** Lipid species correlation of SF–EVs. Combined heatmap (cluster dendrogram) of Lipid–Lipid Spearman correlations between the 50 most abundant lipid species in all EV sample groups. Lipid order was based on Partitioning Around Medoids, also known as K–Medoids, a centroid–based clustering algorithm. On top of the figure is the cluster dendrogram. Below is the group distribution, the relative lipid intensity of each species, and the heatmap. Under the heatmap, the degree of saturation, the lipid class of each lipid, and the respective annotation of each lipid species are indicated. **C)** Changes in EV lipid classes during OA development. Vertical slices plot of SF–EVs showing the relative molar abundances for individual lipid classes. Abbreviations: LysoPC, (lysophosphatidylcholine); LysoPG, (lysophosphatidylglycerol); LysoPI, (lysophosphatidylinositol); LysoPS, (lysophosphatidylserine); PC, (ester–linked phosphatidylcholine); PC O–, (ether–linked phosphatidylcholine); PE, (ester–linked phosphatidylethanolamine); PE O–, (ether-linked phosphatidylethanolamine); PI, (phosphatidylinositol); PS, (phosphatidylserine); SM, (sphingomyelin).

A Spearman correlation heatmap showed that the EV populations of the three different clinical groups differed in the distribution of their phospholipid composition (Fig. 2B). The heatmap was split into three sections (slices and clusters) based on Partitioning Around Medoids (PAM) clustering. For the first slice, the predominant lipid classes were phosphatidylserine (PS), ester–linked phosphatidyl-choline (PC), ester–linked phosphatidylethanolamine (PE) and ether–linked phosphatidylethanolamine (PE–O), which account for half of the lipid distribution of EVs in the healthy joints but for less in both OA groups. The second slice included other members of the PS and PC classes, and the phosphatidic acid (PA), lysophosphatidylcholine (LysoPC), and phosphatidylinositol (PI) lipid classes. There were no clearly identified clusters in this slice. The third slice consisted solely of sphingomyelin (SM), and the distribution was one–third per group; thus, the OA–derived EVs had a higher presence than the EVs from healthy joints. These results showed a subtle variance among SF–EVs from the healthy and the mild and severe OA phenotypes.

#### 3.2.2. Differences in lipid class composition of synovial fluid extracellular vesicles are related to osteoarthritis progression

Having established a difference between the SF–EVs from healthy joints compared to mild OA and severe OA SF–EVs, we proceeded to analyse in more detail how the lipid classes were distributed in the respective groups (Fig. 2C, Suppl. Fig. 2). The most abundant phospholipid classes in all three clinical groups were SM (20–40%), PC (25–40%), PS (12–16%), PE O– (8–12%) and PE (5– 10%) (Fig. 2C). However, a relative increase of SM was observed in the OA groups (healthy 19.9%, mild OA 35.5% and severe OA 37.5%), while the amounts of PC, PI and PS relatively decreased in OA groups which was most pronounced in the severe OA group (healthy: PC 38.3%, PI 1.73% and PS 16.0%; mild OA: PC 31.8%, PI 1.27% and PS 13.35%; severe OA group: PC 26.3%, PI 0.59% and PS 11.8%). Additionally, compared to healthy SF–EVs, ether–linked phosphatidylcholine (PC O–) and PA classes demonstrated a relative rise in mild OA SF–EVs (healthy: PC O– 1.71% and PA 1.02% mild OA: PC O– 1.90% and PA 1.34%). However, the levels declined in severe OA–derived SF–EVs even more than the baseline levels in healthy joint derived EVs (severe OA: PC O– 1.19% and PA 0.55%). Inversely, both PE types (ester–linked and ether–linked) showed a reduction in the mild OA–derived EVs compared to the healthy joint derived EVs (healthy: PE 8.06% and PE O–11.8%; mild OA: PE 5.72% and PE O– 8.17%), while there was an increment in EVs isolated from the severe OA group (Severe OA: PE 10.2% and PE O– 10.2%) with the ester–linked PE class level even higher than in EVs derived from healthy joints.

Despite variations in the total lipid classes with respect to the whole phospholipidome, the individual lipid species contributing to the lipid classes were similarly distributed throughout the clinical groups following normalisation within each class (Suppl. Fig. 2). Thus, the observed shifts in lipid classes cannot be directly attributed to changes in individual lipid species. Overall, these findings demonstrate that the phospholipidome is gradually transformed as OA develops.

### 3.3. Proteomic Analysis

#### 3.3.1. Principal component analysis of proteomics demonstrates variable protein distribution according to osteoarthritic phenotype

Unsupervised multivariate analysis using PCA was conducted on the proteome of all samples exploring the variability between SF–EVs derived from healthy joints, mild OA and severe OA. A total of 5774 unique peptides were identified, translating to 290 proteins with no missing values. Missing values as such were imputed (using impute 2,1,1) using the following method: For the 7 healthy samples, up to 2 missing values were imputed by inserting the mean of the healthy values for that particular protein. Similarly, for the 4 mild OA and 3 severe OA samples, up to 1 missing value was imputed by inserting the mean of the mild OA or severe OA values for that particular protein, resulting in a total of 598 proteins identified and quantified across all samples and used for statistical analysis (Supplementary Figure 1). The first two components (Fig. 3A) reduce the total variation of all the individual data points by 36.4%.

**Figure 3:**
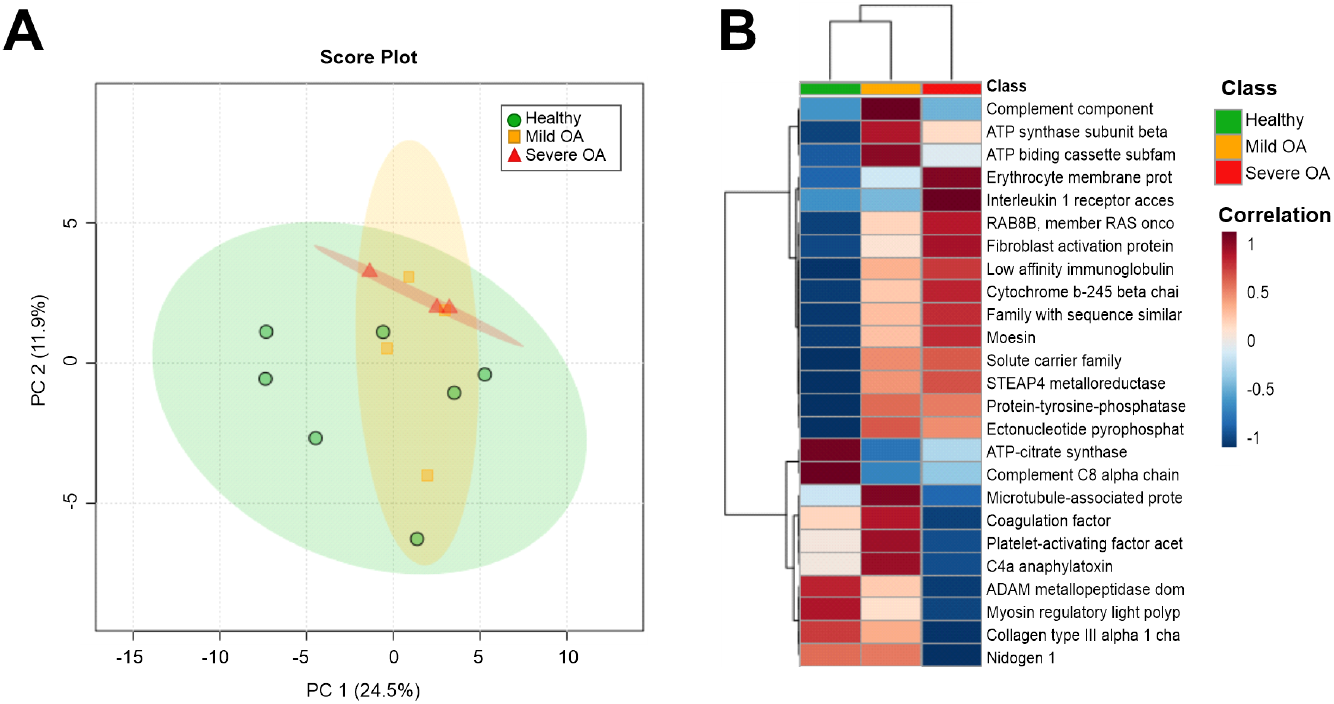
| Proteomic profile of equine SF–EVs derived from healthy joints and from joints with mild and severe OA **A)** Unsupervised multivariate analysis using principal component analysis. The first two principal components were plotted, accounting for 36.4% of the variance. SF–EV samples were plotted based on acquired SWATH–MS data, after PQN normalisation and log transformation. Each plotted point represents a pooled SF–EV sample comprised of three biological replicates, which are colour–coded by OA severity, with severe OA in red, mild OA in orange, and healthy in green. **B)** Heatmap demonstrating average protein intensities between SF–EV healthy (green), mild OA (orange) and severe OA (red) phenotypes. Protein intensities were transformed and are displayed as colours ranging from red to blue. Both rows and columns are clustered using the Ward method, and distance was calculated using Pearson Distance.

#### 3.3.2. Differentially expressed proteins identified across osteoarthritic phenotypes

Using ANOVA, 40 proteins were identified as being significantly differentially expressed (p<0.05) prior to false discovery rate (FDR) adjustment across all experimental groups. Raw values were used due to this being an exploratory study, whereby multiple testing correction methods can fail to identify statistically significant values due to stringent thresholds [46]. Table 1 demonstrates the top 25 differentially expressed proteins and their respective fold change expression. It was revealed that microtubule–associated protein (p=0.006) was present at higher levels in mild OA compared to the severe form of the disease. Further proteins with an increased expression in severe OA compared with the group with healthy joints and mild OA were fibroblast activation protein alpha (p=0.03) and Interleukin 1 receptor accessory protein (p=0.02). Conversely, platelet–activating factor acetylhydrolase IB subunit alpha (p=0.004) exhibited increased expression in mild OA but was decreased in severe OA, as shown in Table 1. Other significant (p<0.05) proteins attributed to EVs that were identified in our dataset included RAB GTPases, such as RAB GDP dissociation inhibitor (p=0.03) and RAB8 (p=0.004). Overall, a change in the proteome was observed in response to an altered OA phenotype, with significant proteins attributed to pathways known for propagating OA disease development within the joint.

**Table 1.**
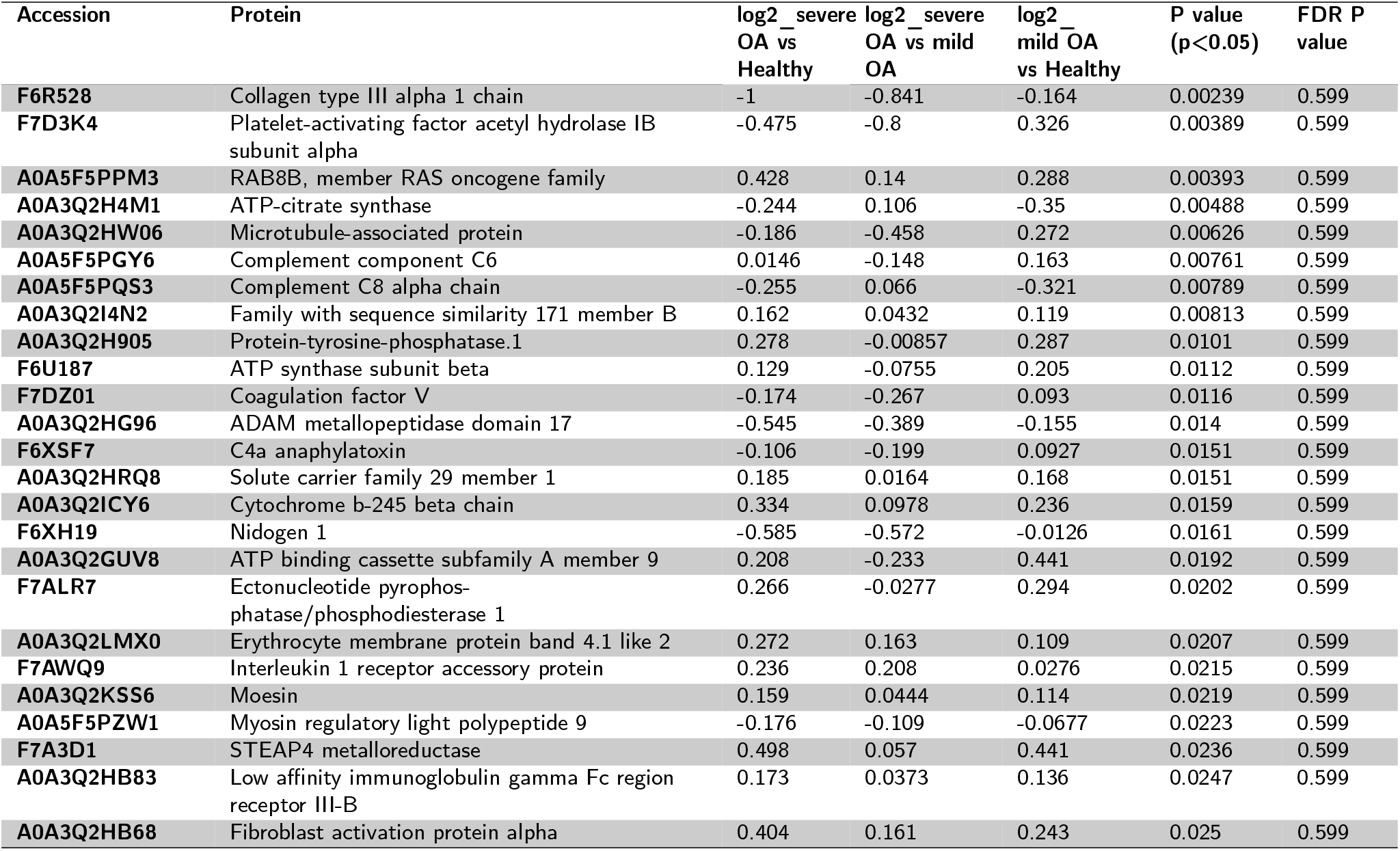
Top 25 differentially expressed (p<0.05) proteins across SF–EV samples derived from healthy joints and joints with mild OA and severe OA following analysis of variance (ANOVA) and heatmap analysis.

#### 3.3.3. A stepwise change in protein expression correlates to osteoarthritis severity

Heatmap analysis was performed on SF–EV samples from healthy joints and mild and severe OA using the Ward clustering method and Pearson distance [42], with selected proteins identified following ANOVA, visualising the top 25 most differential proteins. A stepwise expression change of 10 proteins was observed as OA severity increased, i.e., RAB8B (p=0.0039), moesin (p=0.02), fibroblast activation protein alpha (p=0.03), cytochrome b–245 beta chain (p=0.016), family with sequence similarity 171 (p=0.008), solute carrier family 29 member 1 (p=0.02), STEAP4 metalloreductase (p=0.02), protein tyrosine phosphatase 1 (p=0.01), and ectonucleotide pyrophosphatase (p=0.02) (Fig 3B and Table 1). It was found that 4 EV–associated proteins had an inverse correlation with an increased expression in mild OA but lower expression in EVs derived from severe OA patients (i.e., microtubule–associated protein (p=0.006), coagulation factor V (p=0.01), platelet–activating factor (p=0.004) and c4a anaphylatoxin (p=0.01) (Fig. 3B and Table 1).

#### 3.3.4. Functional enrichment analysis of the synovial fluid–derived extracellular vesicles proteome using IPA highlights dysregulation in pathways associated with cartilage homeostasis and an inflammatory phenotype

Functional enrichment analysis was performed in order to provide biological meaning to the identified and quantified proteome. In both mild OA and severe OA groups, the top canonical pathways were identified using the ingenuity knowledge base library and accounting for protein p-value and log2 fold change of those 40 proteins identified as differentially following ANOVA analysis, 558 of the total 598 proteins were recognised and used in IPA analysis. It was found that signalling by Rho family GTPases (p=1.24×10^−19^), RAC signalling (p=3×10^−21^), actin cytoskeleton signalling (p=3.88×10^−32^), acute phase response signalling (p=5.69×10^−30^), complement system activation (p=1.67×10^−25^), integrin signalling (p=2.72×10^−24^) and clathrin-mediated endocytosis (p=1.30 ×10^−24^), were all significant to OA pathology when considering EV cargo, as shown in Fig. 4 A, B, and C. Additionally, significant diseases and functions in both severe and mild OA included organismal injury (p= 3.98×10^−14^ – 1.58×10^−49^ (confidence interval), inflammatory disease (p= 1.21×10^−14^ – 3.19×10^−39^), and inflammatory response (p= 3.48×10^−14^ – 4.4×10^−38^). Molecular functions that were upregulated in OA groups compared to samples from healthy joints included cell–to–cell signalling and cell movement (p=4.15×10^−14^ – 3.26×10^−63^). Disease and molecular functions unique to the severe OA phenotype were also identified and included an activated state for the following functions: binding of macrophages (p=1.22×10^−16^), interaction of macrophages (p= 1.75 ×10^−17^), immune response of cells (p= 3.20 ×10^−25^), cell movement of antigen–binding cells (p= 1.29 × 10^−20^) and cellular movement of macrophages (p= 1.16 ×10^−18^), as shown in Suppl. Table 4. These data potentially demonstrate a molecular shift in pathophysiology with OA severity with respect to inflammation and immune system involvement.

**Figure 4:**
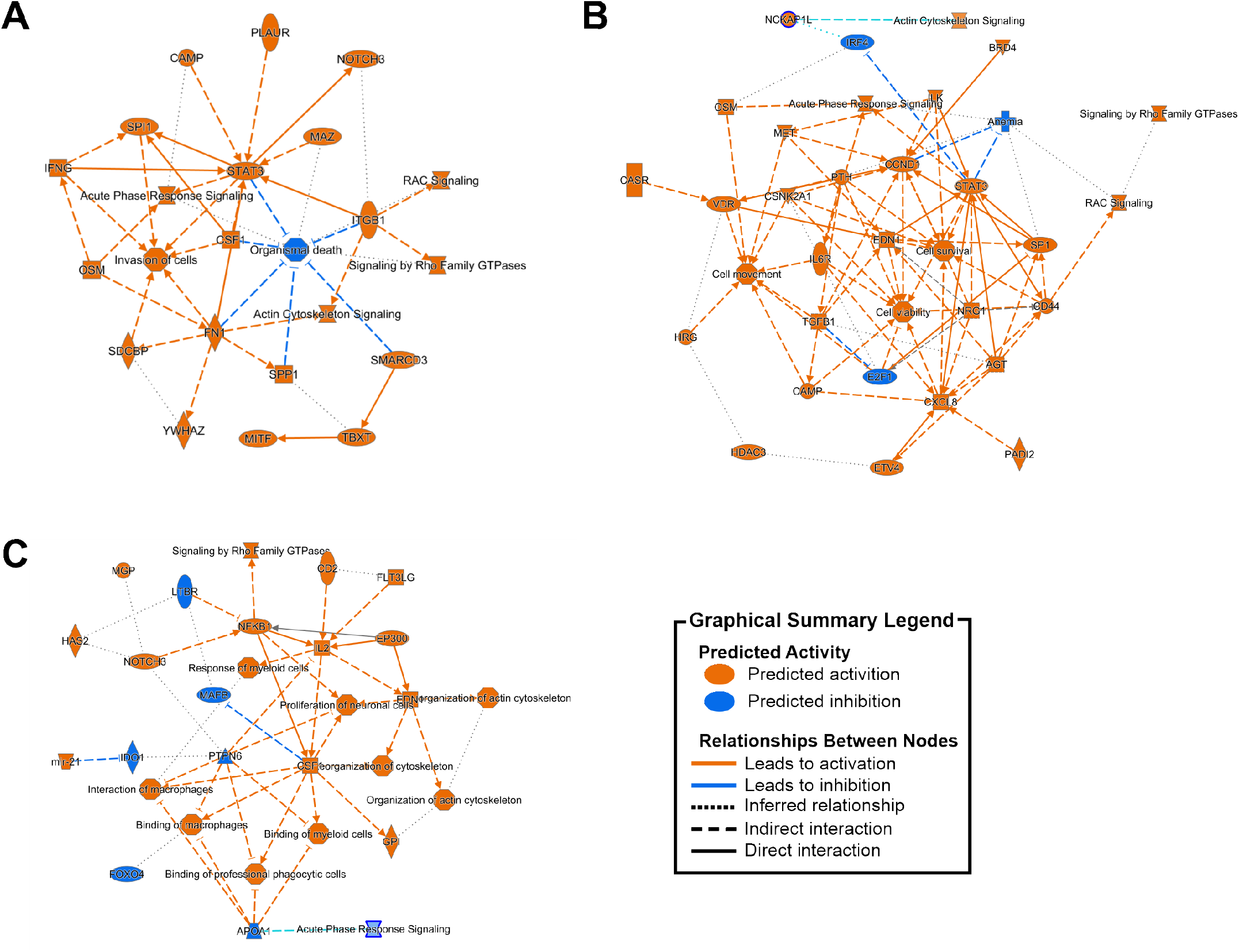
Ingenuity Pathway Analysis networks providing an overview of related molecular mechanisms. Analysis was conducted using the ingenuity knowledge base library and accounting for protein p–value and log2 fold change of the 40 proteins identified as differentially following ANOVA analysis. In each figure, orange denotes upregulation or activation, and blue downregulation or inhibition. **A)** Mild OA compared to healthy, **B)** Severe OA compared to healthy, **C)** Severe OA compared with mild OA.

### 3.4. Multi–Omic Integration

#### 3.4.1. Proteomics and lipidomics data integration demonstrates a high correlation between proteins and phospholipids in synovial fluid extracellular vesicles

Integration of the proteome and phospholipidome datasets was performed to determine if biologically feasible correlates could be established; and thus, identify candidate composite protein–lipid biomarkers. An unsupervised approach was selected to integrate the dataset, consisting of a PCA assessment followed by sparse Partial Least Squares (sPLS2) regression which was tuned by cross–validation.

The initial exploratory analysis employing PCA was undertaken to recognise how the individual proteomic and lipidomic datasets behaved under the same normalisation conditions and to determine the optimal data integration model (Suppl. Fig. 3A, 3B). The omics datasets were normalised by the summed intensity of the sample, followed by centring and scaling of the data, thus subtracting the mean and dividing by the standard deviation. It was observed that clustering of the samples was comparable to the previous PCA (Fig. 2A, Fig. 3A).

Subsequently, to integrate the omics data sets, the unsupervised sPLS2 model was constructed separately for the proteomics and lipidomics data (Suppl. Fig. 3C, 3D). As an unsupervised analysis, the information about the groups (healthy joints, mild OA and severe OA) was not taken into consideration; however, the samples were labelled to understand how they clustered. In Suppl. Fig. 3C and 3D, both sPLS2s project the respective data similarly, with the superior subspace primarily composed of SF–EV samples from healthy joints, the inferior one of mild OA and severe OA SF–EV samples and the top left subsection of overlapping samples from all groups. Afterwards, both sPLS2s were averaged (Fig. 5A). The integrated averaged sPLS2 had a similar structure in components as the individual sPLS2. Figure 5B assesses the degree of agreement between the proteomic and lipidomic datasets by plotting the position of each sample from both sPLS2s in the same space and connecting them with an arrow that indicates at its base the location in the proteomics data set and at the tip the location in the lipidomic data set. Most samples were located relatively close to each other indicating a correlation between the phospholipidome and proteome of SF–EVs.

**Figure 5:**
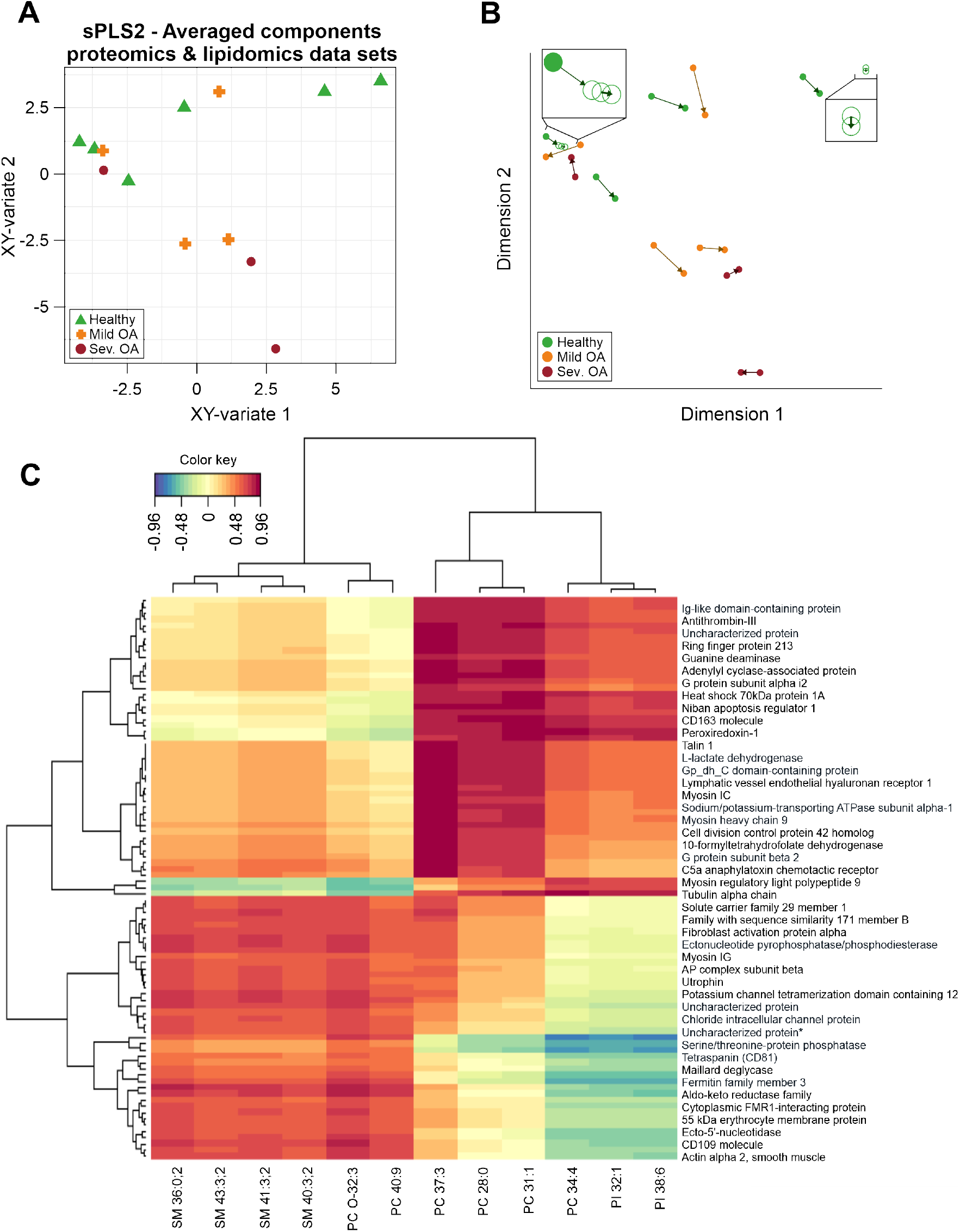
Unsupervised proteomic and lipidomic data integration. Proteomic and lipidomic datasets from SF–EVs derived from healthy joints, mild OA and severe OA were normalised by the sum. **A)** Sparse Partial Least Squares–2 regression (sPLS2) of SF–EV samples projected into the area covered by the averaged components of both datasets. Healthy SF–EVs (green triangle), mild OA SF–EVs (orange cross), and severe OA SF–EVs (Sev. OA; red circle). **B)** Unsupervised multivariate sPLS2 arrow plot from the integration of proteomic and lipidomic data. The base of the arrow shows where a specific sample is in relation to the components of the proteomics dataset, and the tip of the arrow shows where the same sample is located concerning the components of the phospholipidomics dataset. Healthy SF–EVs (green circle), mild OA SF–EVs (orange circle), and severe OA SF–EVs (Sev. OA; red circle). The boxes zoom in on certain samples to better show the arrow direction **C)** Clustered Image Map from the sPLS2 data integration performed on the SF–EV omic datasets. The graphic shows the degree of similarity between the proteomic and lipidomic variables clustered over two dimensions and grouped using the Euclidean distance approach.

This correlation was further explored with a Cluster Image Map (CIM) (Fig. 5C) to examine the connection between the features and components in a broad range, drawing attention to the relevant variables that collectively accounted for the covariance between the two datasets. According to the CIM, the phospholipid variables were divided into three slices that were either positively or negatively related to two main protein clusters. The left slice corresponded to 4 SMs (SM 36:0;2, SM 43:3;2, SM 41:3;2, SM 40:3,2), PCO–32:3, and PC 40:9, which had a positive association with the lower protein cluster. The middle slice, consisting of three PC species (PC 37:3, PC 28:0 and PC 31:1), had an inverse pattern of the cluster, with the upper group depicting the strongest association. Finally, the right slice, comprising the PC 34:4 and the two PI species (PI 32:1 and PI 38:6), had a similar association pattern as the middle one; however, the lower cluster exhibited a negative correlation, while the cluster above was positively correlated to the proteins.

#### 3.4.2. Relevance network for the selection of candidate proteins and phospholipids as composite OA biomarkers

To better comprehend the correlation between the proteins and phospholipids, a relevance network plot was created (Fig. 6). Three substructures could be identified from the network. The larger cluster contained the same lipids as the middle slice from the CIM (Fig. 5C; PC 28:0, PC 31:1 and PC 37:3), with all the correlations depicted being positive. The second substructure consisted of the right–side slice lipids from the CIM (Fig. 5C; PC 34:4, PI 32:1, PI 38:6), with primarily positive correlations to the proteins except to the anion exchange protein. This cluster also overlapped with some of the same proteins as PC 28:0, PC 31:1 and PC 37:3. The third substructure was composed of the lipids from the left–side slice of the CIM (SM 36:0;2, SM 43:3;2, SM 41:3;2, SM 40:3,2, PC O–32:3, PC 40:9). This cluster displayed only positive correlation with the depicted proteins, including the anion exchange protein. Moreover, the proteins that correlated to phospholipids from the relevance network plot (Fig. 6) were found to be associated with pathways such as actin cytoskeleton signalling (p=5.71×10^−7^) and signalling by Rho family GTPases (p=5.89×10^−8^), as shown in Table 2.

**Table 2.** Top 5 canonical pathways identified using IPA, following input of proteins correlated to lipids.

| Canonical Pathways | P-Value | Number of annotated molecules |
| --- | --- | --- |
| Leukocyte extravasation signalling | 1.64E-10 | 9/193 |
| RHOGDI signalling | 5.24E-10 | 9/220 |
| Fcy Receptor-mediated phagocytosis in macrophages and monocytes | 3.52E-08 | 6/94 |
| Signalling by Rho Family GTPases | 5.89E-08 | 8/267 |
| Actin cytoskeleton signalling | 5.71E-07 | 7/244 |

**Figure 6:**
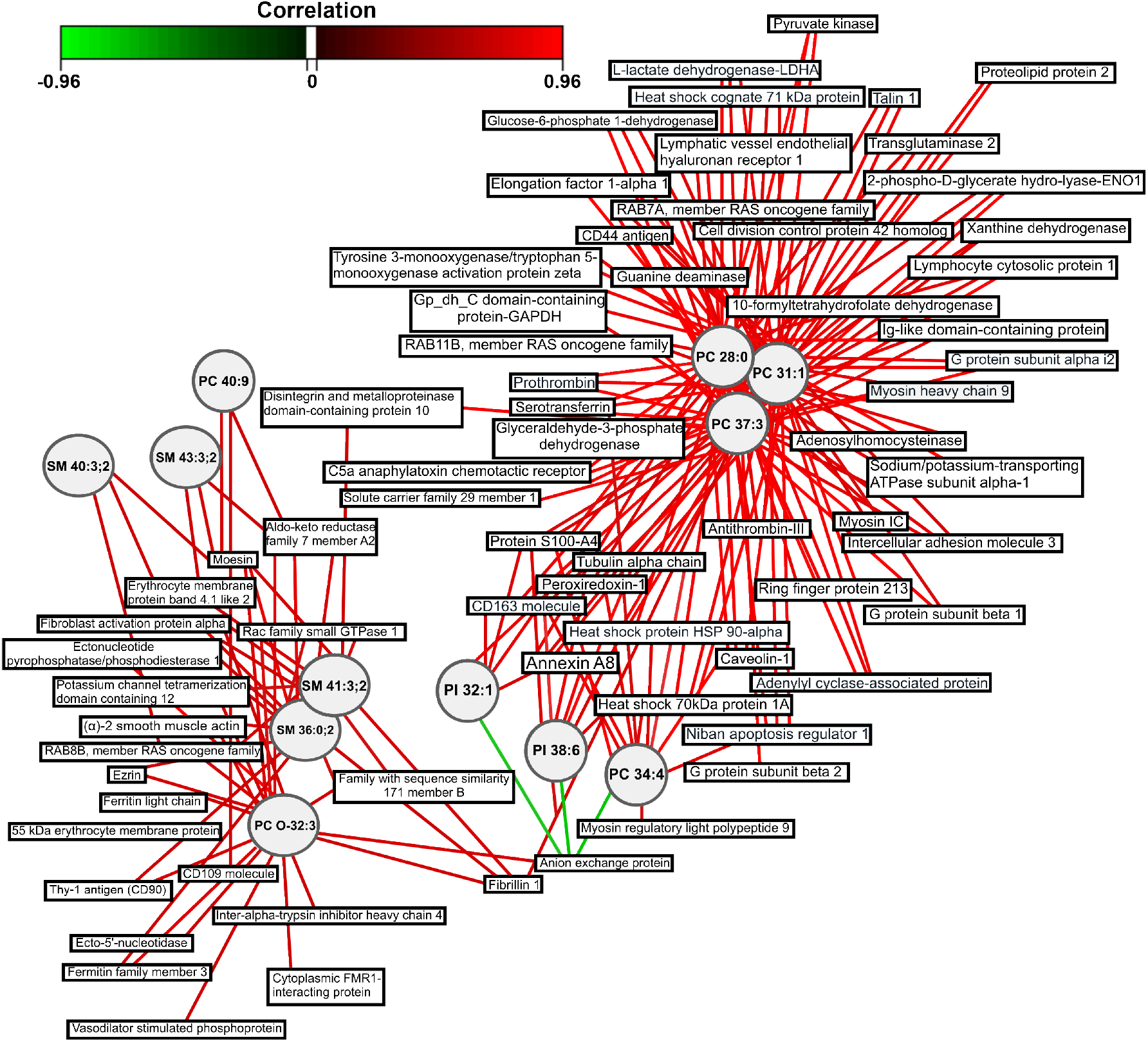
Network representation derived from the sPLS2 analysis of the proteomics and lipidomics integrated data. A relevance network plot with a correlation cutoff of 0.7 was created. Hence, only the variables with a correlation above 0.7 or below –0.7 are shown. The networks are bipartite, and each edge connects a protein (rectangle) to a phospholipid (circle) node based on a similarity matrix. The colour of the lines represents positive (red) or negative (green) correlations.

Since the sPLS2 analysis is an unsupervised approach (i.e., no information regarding the groups is entered in the model), neither the CIM nor the network explained how the SF–EV phospholipids and the correlated proteins relate to the healthy joints, mild OA and severe OA. To determine differences between the clinical groups, all lipids and proteins with a correlation above 0.754 based on the network (Fig. 6) were assessed with a Kruskal–Wallis test (Fig. 7). A significant decrease in PC 34:4 and PI 38:6, and decline in PI 32:1 and the related proteins showed a similar trend in SF–EVs derived from severe OA compared to healthy joints (Fig. 7). Conversely, SM 36:0;2, SM 41:3;2 and PC O–32:3 and the correlated proteins showed a trend to increase with the severity of OA, with significant differences for PC O–32:3, moesin and vasodilator–stimulated phosphoprotein (Fig. 7). A list of potential candidate proteins for composite biomarkers is provided in Suppl. Table 5. Overall, we here show a strategy for composite biomarker discovery based on SF–derived EV–associated phospholipids and proteins and revealed potential candidates that could be explored as composite phospholipid–protein EV biomarkers in OA pathology.

**Figure 7:**
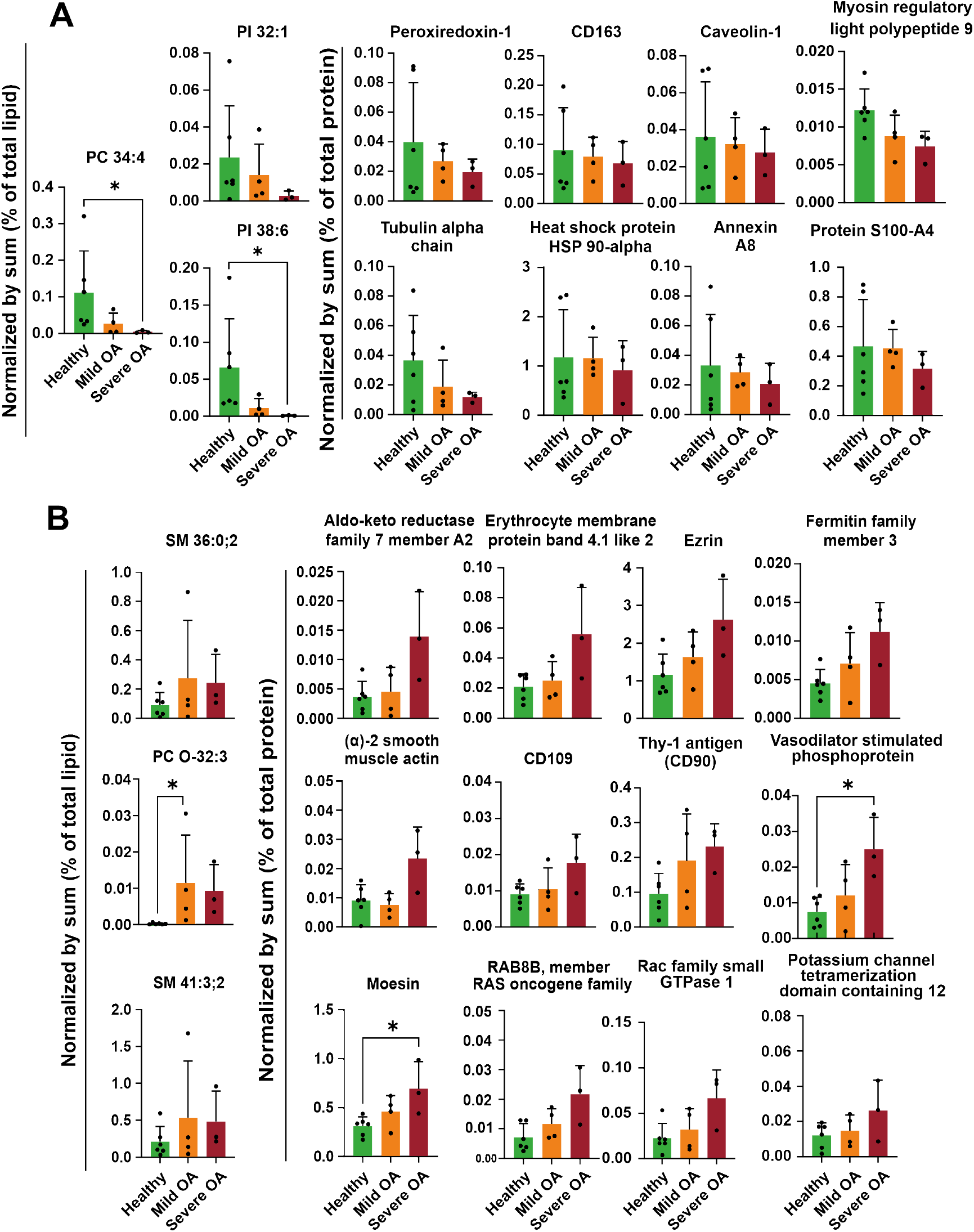
Discovery of potential composite biomarkers for OA. Phospholipids and proteins were normalised by the sum of the total amount of material (i.e., lipid or protein). Healthy (green), mild OA (orange) and severe OA (red). * p < 0.05, Kruskal–Wallis with Dunn’s post hoc test **A)** Phospholipids and proteins that decreased with OA severity, **B)** Phospholipids and proteins that increased with OA severity.

## 4. Discussion

In this exploratory study we designed a workflow for a multi–omics approach based on phospholipidomic and proteomic integration to identify composite SF–derived EV–biomarkers for OA, based on the analysis of EVs isolated from SF of horses with clinically defined OA (mild OA and severe OA), or from healthy equine joints. Hereto we investigated the phospholipidome and proteome of purified SF–EVs and designed a strategy for multi–omics data integration and differential expression analysis. To identify genuine composite EV–biomarkers, we used differential centrifugation followed by sucrose density gradient centrifugation to purify EVs from SF by removing most types of the contaminating lipoproteins and protein aggregates [16; 21]. While the numbers of EVs in SF were unaffected by OA, consistent with other studies [47; 48], the proteomic and phospholipidomic profiles of SF–EVs were correlated to the presence of OA.

We found that OA pathology directly impacted the phospholipidome at the lipid class level, showing gradual changes in several lipid classes associated with disease severity. The relative reduction in PC, PS and PI in mild OA and more drastically in severe OA, could be explained by the relative increase in SM. Since PC and SM are primarily located in the outer layer of the plasma membrane, the increase in SM disrupts the balance and reduces the amount of PC [49]. Similarly, although PS and PI are predominantly found in the inner leaflet of the lipid bilayer, they can also be affected by an increase in SM. Furthermore, we found higher levels of PC compared to PC O–, while the PE and PE O– classes showed an opposite trend. These findings align with the lipidomics findings in the EV field and highlight the importance of ether lipids, especially PE O–, in EV biology, including membrane trafficking and cholesterol regulation [50]. SM, one of the main lipid classes detected in the SF–EVs, plays a crucial role in the plasma membrane composition, cellular proliferation, differentiation, growth, signal transduction, and apoptosis [51]. SMs are instrumental in the formation of lipid rafts enabling the selection of membrane proteins involved in signal transduction and intracellular transport [52]. The notable increase of SMs with OA severity suggests that more lipid raft–like domains may be present in SF–EVs as the OA pathology progresses, facilitating and enhancing the cell–to–cell communication of SF–EVs.

Functional enrichment analysis of the differentially expressed SF–EV proteins identified a range of activated canonical pathways associated with disease phenotype. Specifically, Rho family GTPases, including RAC family small GTPase1, actin, and integrin–associated proteins, were identified as activated in both OA groups (mild and severe) compared to healthy joints. Dysregulation of Rho GTPases has been implicated in rheumatic disorders in humans like rheumatoid arthritis, OA, and psoriatic arthritis, contributing to hypertrophic changes and cartilaginous matrix destruction [53–55]. Rac1, a pro–inflammatory factor, stimulates MMP13 production and upregulates markers of chondrocyte hypertrophy, such as COLX and ADAMTS–5 [54]. Dysregulated activation of Rho GTPases, particularly CDC42, can lead to the degradation of articular chondrocytes through IL–6/STAT3 signalling [56]. The presence of these proteins in SF–EVs from diseased groups suggests their potential role in propagating disease within the joint by carrying cargo that induces phenotypic and metabolic changes.

Integrin signalling was also enhanced in SF–EVs from OA joints (mild and severe), indicated by increased expression of integrin–linked kinase, integrin subunits alpha 1, 5, and 6, and integrin subunits beta 1, 2, and 5. Integrins are adhesion molecules involved in signal transduction and mechanical deformation through cytoskeleton contraction [57]. They play a crucial role in OA pathogenesis by mediating cell–matrix interactions, inflammation promotion, and immune cell recruitment [58]. Our study observed enrichment of these integrin subunits in OA EVs, suggesting their involvement in OA pathogenesis. Functional enrichment analysis also revealed upregulation of disease and molecular functions related to macrophage binding, interaction, and cellular movement in severe OA compared to mild OA. These findings suggest the significant involvement of macrophages in the later stages of OA pathogenesis. Previous studies have identified that an imbalance of macrophage subtypes (M1 and M2), can contribute to the chronic low-grade inflammation associated with OA and is implicated in OA pain mechanisms [59]. Additionally, macrophages play a crucial role in regulating inflammation and are known mediators of OA–related inflammation [60].

Overall, the observed changes in both phospholipid classes and proteins between SF–EVs derived from healthy joints and OA patients and the gradual changes associated with the severity of OA, suggest that these SF–EV parameters may be used as natural composite biomarkers for OA diagnosis and progression. Our multi–omics integration approach, using unsupervised sPLS2 regression and PCA, indeed revealed a remarkably strong similarity in the space distribution induced by the SF–EV phospholipidome and proteome, indicating a strong interrelationship, which is mainly due to a strong correlation between specific phospholipids with a certain set of proteins. Functional enrichment analysis of the proteins from this correlation network revealed several canonical pathways, such as signalling by Rho Family GTPases as previously identified and actin cytoskeleton signalling.

Integration of data revealed potential composite biomarkers consisting of downregulated and upregulated phospholipids and proteins as OA severity progressed. Downregulated proteins and the respective phospholipids were comprised of phospholipids PC 34:4, PI 32:1 and PI 38:6, and proteins such as heat shock protein 90 (HSP90AA1) and CD163. Interestingly, HSP90AA1 has been demonstrated to be down–regulated in blood and cartilage of human patients with OA, and levels correlated with the risk incidence of OA [61], while CD163, a transmembrane protein of M2 macrophages [62], was shown to decline as OA progressed in this study. It has been suggested that the inability of macrophages to transform from M1 to M2 might contribute to the onset and development of OA [63].

Among the upregulated proteins, several structural proteins were detected, including α–2 smooth muscle actin, erythrocyte membrane protein band 4.1–like 2 (EPB41L2), ezrin, and moesin. These proteins likely indicate changes in diseased joint tissues, which were reflected in the structural protein composition of SF–EVs. α–smooth muscle actin is known to be expressed in fibroblast–like synoviocytes (FLSs) undergoing a change to a myofibroblast–like phenotype in the presence of transforming growth factor *β* (TGF*β*), linked to OA pathogenesis [64], as well as to colocalise with fibronectin, which is associated with inflammation in OA [65]. Ezrin, moesin, and EPB41L2 activation promote enhanced proliferation and formation of fibrillated OA cartilage by blocking cell–cell contact inhibition in chondrocytes [66]. Ezrin has also been connected to the RhoGTPase signalling pathway in OA synovial fluid [66]. Additionally, CD90 and CD109 transmembrane proteins, upregulated as OA progresses, regulate the pathological response in rheumatoid arthritis (RA) fibroblast–like synoviocytes, driving inflammation and fibrosis [67; 68]. The upregulated proteins were associated with phospholipids SM 36:0;2, SM 41:3;2, and PC O–32:3. The combinations of these proteins and phospholipids could potentially serve as candidate composite SF–EV biomarkers for OA onset and progression.

The inherent constraints of this exploratory study, such as relatively small clinical sample size and large volume of SF required, non–conformity of radiological and clinical parameters for OA severity assignment and lag in the development of analytical tools when comparing mass spectrometry proteomics and lipidomics pipelines, are important to overcome in future studies. Nonetheless, the approach of our exploratory study in equine OA highlights the potential for identifying important molecular mechanisms of OA and aims to serve as a framework for the discovery of SF–derived EV–based composite biomarkers having the potential to inform disease severity and enable targeted disease management in the future.

## 5. Funding

Author **L.V**. received funding from the EU’s H2020 research and innovation program under Marie S. Curie CO-FUND RESCUE grant agreement No 801540. **E.C**. is a self–funded PhD student from the University of Liverpool and acquired funding from the EUCost initiative (ExRNA path) cost action CA20110. **E.L.A**. received funding from the EU’s H2020 research and innovation programme under the Marie Skłodowska–Curie grant agreement No 722148 (TRAIN–EV). **M.P**. and **E.C**. were also supported by the Medical Research Council (MRC) and Versus Arthritis as part of the MRC Versus Arthritis Center for Integrated Research into Musculoskeletal Aging (CIMA). **M.P** was supported by a Horserace Betting Levy Board grant (Prj794).

## 6. Acknoledgements

We thank Jeroen Jansen for running the MS for the lipidomics experiment. The authors acknowledge the use of the CDSS Bioanalytical Facility provided by Liverpool Shared Research Facilities, Faculty of Health and Life Sciences, University of Liverpool.

## CRediT authorship contribution statement

**Emily J Clarke:** Conceptualisation, Formal analysis, Investigation, Project administration, Validation, Visualisation, Writing - Original Draft, Writing – Review & Editing. **Laura Varela:** Conceptualisation, Formal analysis, Investigation, Project administration, Validation, Visualisation, Writing – Original Draft, Writing – Review & Editing. **Rosalind E Jenkins:** Data Curation, Formal analysis, Investigation, Validation, Writing – Review & Editing. **Estefanía Lozano**–**Andrés:** Investigation, Writing – Review & Editing. **Anna Cywińska:** Resources, Writing – Review & Editing. **Maciej Przewozny:** Resources. **P. René van Weeren:** Funding acquisition, Project administration, Supervision, Writing – Review & Editing. **Chris H.A. van de Lest:** Conceptualisation, Data Curation, Formal analysis, Validation, Supervision, Software, Writing – Review & Editing. **Mandy Peffers:** Conceptualisation, Project administration, Supervision, Writing – Review & Editing. **Marca H.M. Wauben:** Conceptualisation, Project administration, Supervision, Writing – Review & Editing.

## Notes

https://www.uu.nl/en/research/yoda

https://www.ebi.ac.uk/pride/

### Competing Interest Statement

The authors have declared no competing interest.

