## Supplementary Material for "Proteome and phospholipidome interrelationship of synovial fluid-derived extracellular vesicles in equine osteoarthritis: An exploratory ‘multi-omics’ study to identify composite biomarkers"

**Table 1:** Radiographic criteria used for the classification of the healthy, mild and severe OA phenotypes. Arrows indicate osteophyte formation and subchondral osteolysis, and R and L denote right or left limb.

| Group | Description | Radiograph |
|---|---|---|
| Healthy | No lesions | 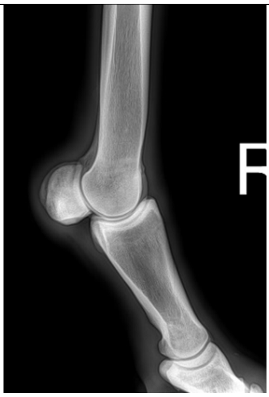 |
| Mild OA | Small osteophyte and subchondral osteolysis | 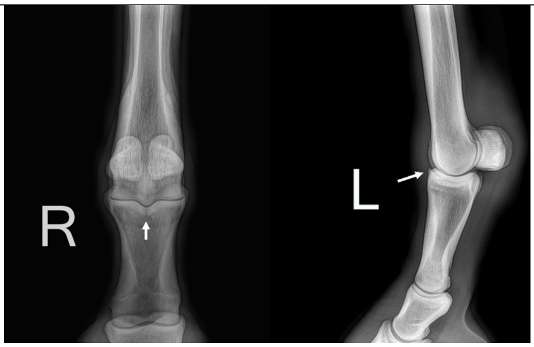 |
| Severe OA | Bone deformation, narrowing of joint space, subchondral bone sclerosis, osteophytes | 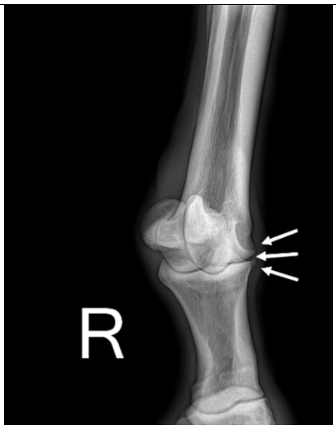 |

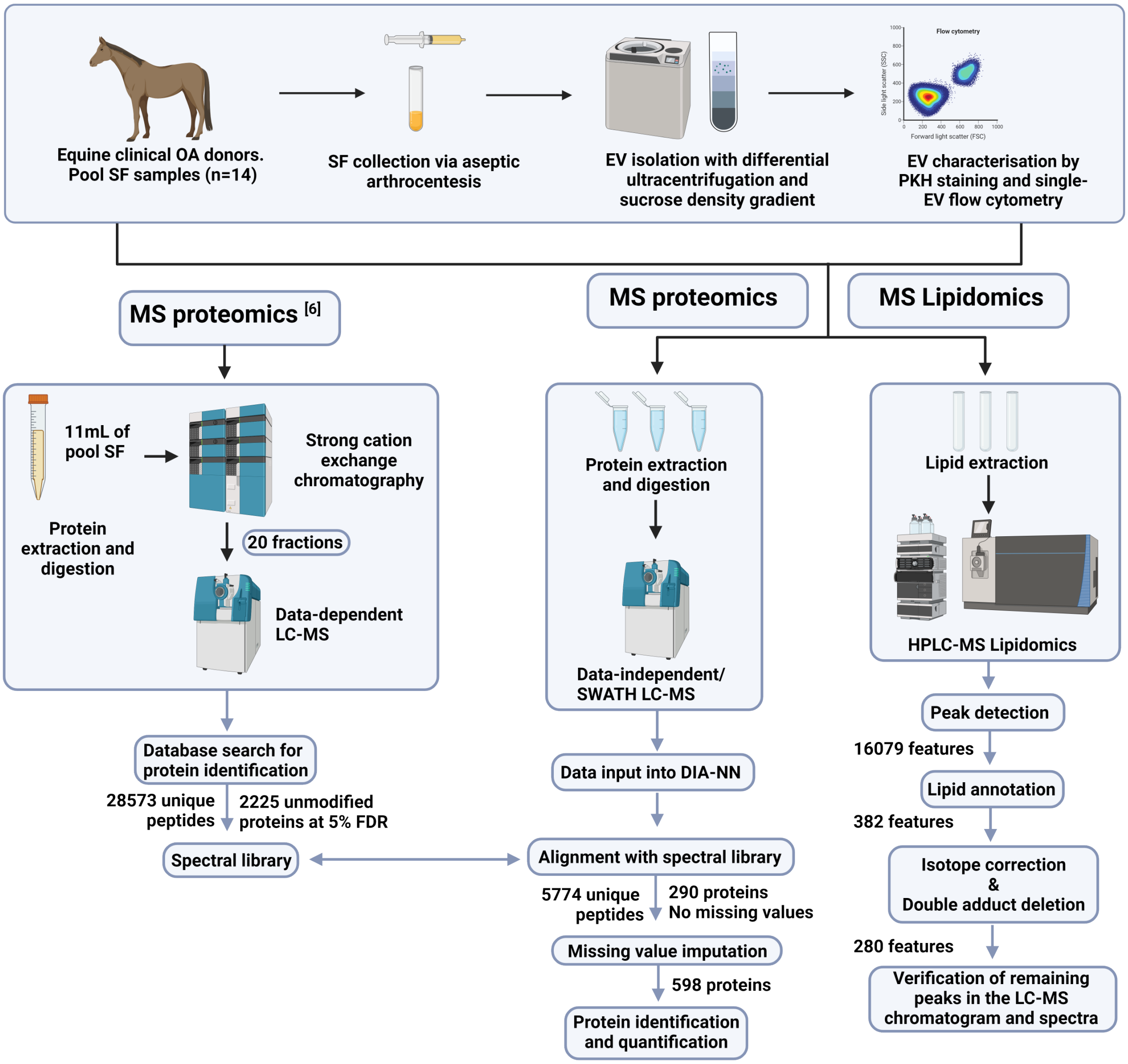


**Suppl. Fig. 1 : Workflow for sample processing.** OA was characterised following clinical and radiographic examinations. SF was collected by sterile arthrocentesis and spun to create cell-free SF. Forty-two donors were used to create 14 samples consisting of a pool of three unique biological samples with a volume of 5mL (Healthy n=7, mild OA n= 4, and severe OA n= 3). EVs were isolated from cell-free SF following differential ultracentrifugation with a sucrose density gradient [1]. EVs were stained using PKH and subsequently characterised following quantitative single EV-based high-resolution flow cytometry [2; 3]. The EV lipidome was probed following a chloroform and methanol lipid extraction [4], and mass spectrometry lipidomics was performed using a Fusion Orbitrap MS passing through a heated electrospray ionisation [5]. The peaks were detected based on retention time, exact m/z-ratio, and, if present in at least 3 out of 13 samples (Healthy n=6, OA n=4, Advanced OA n=3). The features were annotated using an in-silico phospholipid database. The features were also selected to account for isotope distribution and adducts. The EV proteome was extracted using a urea lysis buffer and proteins were subsequently digested on hydrophilic and hydrophobic magnetic carboxylate SpeedBeads, using trypsin/lyseC [6]. A data-independent proteomic approach was utilised in the form of Sequential Windowed Acquisition of all theoretical fragments (SWATH), and analysis was performed using a Triple TOF 6600. Statistical analysis was performed on respective datasets independently with Rstudio or Metaboanalys.

**Table 2:** Author Checklist: MIFlowCyt-Compliant Items.

| Requirement | Requested Information |
|---|---|
| 1.1. Purpose | To investigate extracellular vesicle biomarker discovery in synovial fluid from healthy, mild and severe osteoarthritis (OA) equine samples by using a combined proteomics and lipidomics approach |
| 1.2. Keywords | Osteoarthritis, synovial fluid, proteomics, lipidomics, equine, extracellular vesicles |
| 1.3. Experiment variables | Healthy versus clinically relevant equine OA |
| 1.4. Organisation name and address | Division Cell Biology, Metabolism & Cancer<br><br>Department of Biomolecular Health Sciences<br><br>Faculty of Veterinary Sciences<br><br>Utrecht University<br><br>Yalelaan 2, 3584 CM<br><br>Utrecht, The Netherlands |
| 1.5. Primary contact name and email address | Prof. Dr. M.H.M. Wauben<br><br> |
| 1.6. Date or time period of experiment | October 2022 |
| 1.7. Conclusions | The proteome and lipidome of SF-EVs are primarily positively correlated with activation of pathways related to chondrocyte dysregulation and inflammation. A series of phospholipids and proteins were proposed as alternatives for combined biomarker discovery, for example, PC O-32:3 and CD109. |
| 1.8. Quality control measures | Procedural control for fluorescent stainings<br><br>Serial dilutions<br><br>Detergent treatment control |
| 2.1.1.1. (2.1.2.1., 2.1.3.1.) Sample description | Synovial fluid from healthy equine joints, joints with diagnosed mild OA or severe OA |
| 2.1.1.2. Biological sample source description | Synovial fluid |
| 2.1.1.3. Biological sample source organism description | Horse (Equus caballus) |
| 2.1.2.2. Environmental sample location | |
| 2.3. Sample treatment description | For detergent treatment control samples were treated with 0.1% (v/v) triton X-100 (SERVA Electrophoresis GmbH, Heidelberg, Germany) final concentration for 30 seconds before reanalysis. |
| 2.4. Fluorescence reagent(s) description | PKH67 (Sigma-Aldrich) |
| 3.1. Instrument manufacturer | Becton Dickinson |
| 3.2. Instrument model | BD Influx™ optimised instrument for detection of sub-micron sized particles as described previously. |
| 3.3. Instrument configuration and settings | BD Influx optimised to measure small particles. All configuration details can be found in previous publications [2; 3]. Briefly, samples were measured at a constant flow rate for 30 seconds using a fluorescence threshold on the 488 nm laser. The threshold level was set to detect 10-20 events per second when measuring a buffer control. |

| | |
|---|---|
| 4.1. List-mode data files | All data files, including the quality control measure from 1.8, are available upon request. |
| 4.2. Compensation description | No compensation was required due to instrument configuration. |
| 4.3. Data transformation details | No data transformation was applied. |
| 4.4.1. Gate description | No gates were applied |
| 4.4.2. Gate statistics | The number of total events recorded in 30 seconds measurements are shown in the dot plots without any background correction. |
| 4.4.3. Gate boundaries | N/A |

**Table 3:** MIFlowCyt-EV framework.

| | |
|---|---|
| 1.1 Preanalytical variables conforming to MISEV guidelines | Yes, all relevant data has been submitted to EV-TRACK for transparent reporting and centralising knowledge in extracellular vesicle research (EV-TRACK ID: EV230607). |
| 1.2 Experimental design according to MIFlowCyt guidelines | Yes, MIFlowCyt checklist can be found as part of the supporting information of this manuscript in Suppl. Table 2. |
| 2.1 Sample staining details | Yes, described in Materials and Methods. |
| 2.2 Sample washing details | Yes, described in Materials and Methods. |
| 2.3 Sample dilution details | Yes, described in Materials and Methods. |
| 3.1 Buffer-only controls | Yes, relevant buffer controls were measured |
| 3.2 Buffer with reagent controls | Yes. Data available upon request |
| 3.3 Unstained controls | N/A |
| 3.4 Isotype controls | N/A |
| 3.5 Single-stained controls | N/A |
| 3.6 Procedural controls | Yes. Available upon request |
| 3.7 Serial dilutions | Yes, serial dilutions were performed in previous characterisation experiments to determine the ideal dilution used in this study. Data available upon request |
| 3.8 Detergent-treated controls | Yes, sensitivity to triton X-100 was determined in previous characterisation experiments. Data available upon request |
| 4.1 Trigger channel(s) and threshold(s) | The trigger channel used was on the fluorescent signal collected from the 488 nm laser (530/40 bandpass filter). The threshold level was set at 0.62, allowing an event rate of <20 events/second in a clean PBS sample. Additional details can be found in Materials and Methods. |
| 4.2 Flow rate / volumetric quantification | Yes, low flow rate was kept constant and was measured for quantification purposes. The flow rate was estimated at 12.8 μL/min. |
| 4.3 Fluorescence calibration | N/A |
| 4.4 Scatter calibration | N/A |
| 5.1 EV diameter/surface area/volume approximation | N/A |
| 5.2 EV refractive index approximation | N/A |
| 5.3 EV epitope number approximation | N/A |
| 6.1 Completion of MIFlowCyt checklist | Yes, see Suppl. Table 2 |
| 6.2 Calibrated channel detection range | As shown in previous publications, equivalent to 100 FITC MESF. See description in Arkesteijn GJA et al.[3] |
| 6.3 EV number/concentration | Yes, see Figure 1. |
| 6.4 EV brightness | N/A |
| 7.1 Sharing of data to a public repository | Yes, all experimental details about the biological sample preparation can be found in EV-TRACK. All data files are available upon request. |

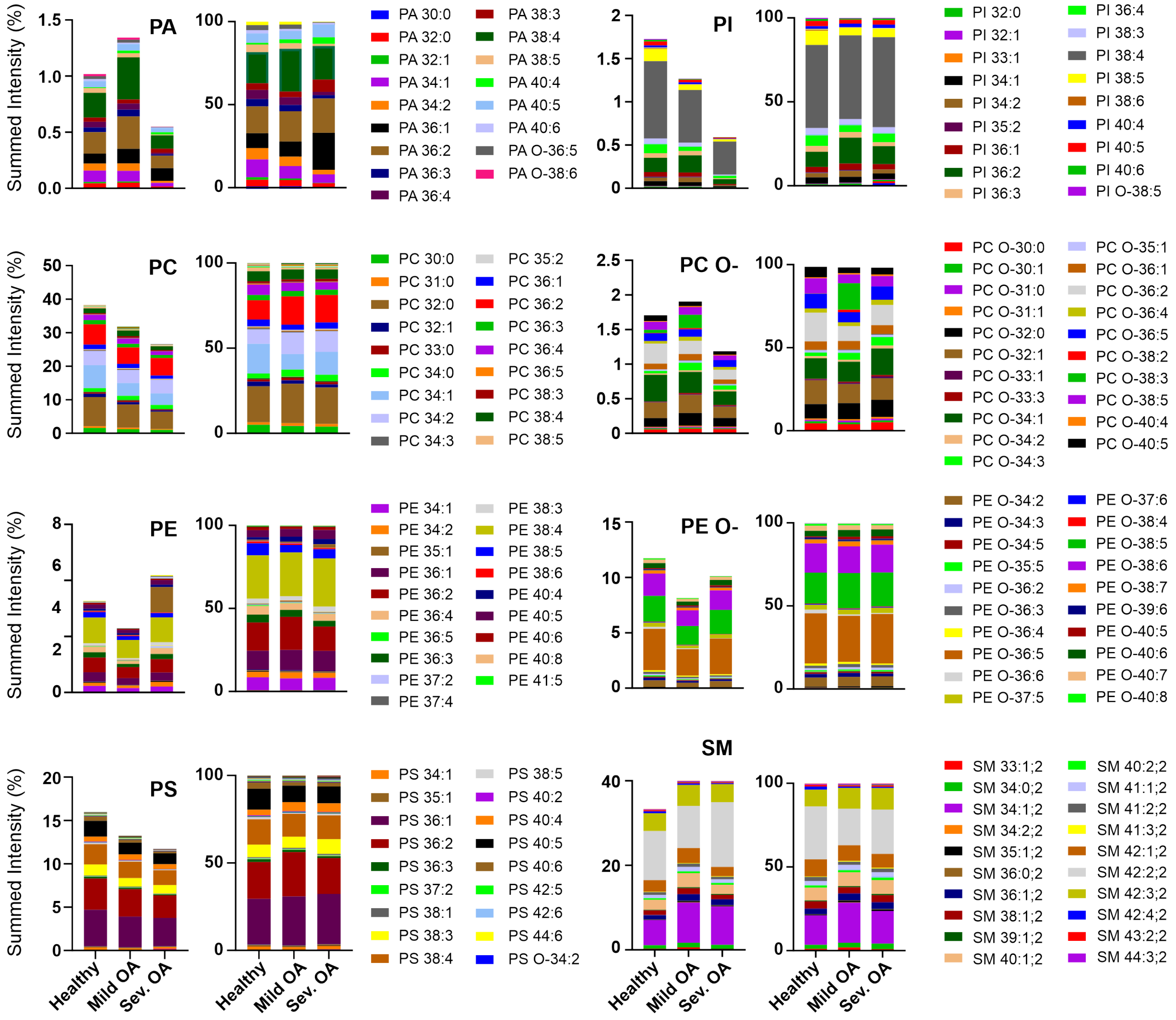


**Suppl. Fig. 2 : Composition of individual lipid species in PA, PI, PC (O-), PE(O-), PS, and SM classes.** Lipid composition of the most abundant lipid classes of SF-EVs from healthy, mild OA, or severe OA horse patients. The left stacked bars graph in each class shows the lipid species' amount in the overall lipidome. The immediately right stacked bars graph displays the normalised amount of lipid species in each class. Samples were normalised for each class and SF-EV group by expressing the lipid intensity as a fraction of the sum of lipid intensities. Lipids were obtained from 100,000g purified SF-EVs with sucrose density gradients. Healthy samples (n=6), mild OA (n=4), severe OA (n=3). Abbreviations: PA (phosphatidic acid), PC (ester-linked phosphatidylcholine), PC O- (ether-linked phosphatidylcholine), PE (ester-linked phosphatidylethanolamine), PE O- (ether-linked phosphatidylethanolamine), PI (phosphatidylinositol), PS (phosphatidylserine), SM (sphingomyelin).

**Table 4:** Significantly differentially expressed pathological functions, as identified by ingenuity pathway analysis for each comparison.

| | Disease or Function Annotation | p-value | Predicted Activation State |
|---|---|---|---|
| Severe OA Vs mild OA | Binding of myeloid cells | 1.76E-40 | Increased in Severe OA |
| | Binding of professional phagocytic cells | 8.25E-35 | Increased in Severe OA |
| | Interaction of phagocytes | 1.31E-34 | Increased in Severe OA |
| | Adhesion of myeloid cells | 3.9E-33 | Increased in Severe OA |
| | Organisation of actin cytoskeleton | 1.34E-31 | Increased in Severe OA |
| | Cell movement of granulocytes | 2.09E-31 | Increased in Severe OA |
| | Phagocytosis | 3E-29 | Increased in Severe OA |
| | Phagocytosis of cells | 1.67E-27 | Increased in Severe OA |
| | Internalisation of cells | 1.42E-25 | Increased in Severe OA |
| | Immune response of cells | 3.2E-25 | Increased in Severe OA |
| | Migration of myeloid cells | 1.56E-23 | Increased in Severe OA |
| | Reorganisation of cytoskeleton | 4.3E-23 | Increased in Severe OA |
| | Transmigration of leukocytes | 2.34E-21 | Increased in Severe OA |
| | Migration of granulocytes | 8.57E-21 | Increased in Severe OA |
| | Cell movement of antigen presenting cells | 1.29E-20 | Increased in Severe OA |
| | Binding of antigen presenting cells | 1.56E-20 | Increased in Severe OA |
| | Cell-cell contact | 2.66E-19 | Increased in Severe OA |
| | Cell movement of macrophages | 1.16E-18 | Increased in Severe OA |
| | Recruitment of cells | 2.14E-18 | Increased in Severe OA |
| | Response of myeloid cells | 3.8E-18 | Increased in Severe OA |
| | Interaction of macrophages | 1.75E-17 | Increased in Severe OA |
| | Immune response of myeloid cells | 8.69E-17 | Increased in Severe OA |
| | Binding of macrophages | 1.22E-16 | Increased in Severe OA |
| | Cell movement | 3.26E-63 | Increased in Severe OA |
| Severe OA vs healthy | Migration of cells | 4.29E-54 | Increased in Severe OA |
| | Adhesion of immune cells | 3.01E-47 | Increased in Severe OA |
| | Endocytosis | 1.36E-44 | Increased in Severe OA |
| | Engulfment of cells | 9.6E-44 | Increased in Severe OA |
| | Cell movement of blood cells | 1.98E-42 | Increased in Severe OA |
| | Leukocyte migration | 3.23E-42 | Increased in Severe OA |
| | Cell movement of leukocytes | 1.22E-40 | Increased in Severe OA |
| | Binding of myeloid cells | 1.76E-40 | Increased in Severe OA |
| | Cell movement of phagocytes | 4.4E-38 | Increased in Severe OA |
| | Cell movement of myeloid cells | 2.1E-36 | Increased in Severe OA |
| | Vasculogenesis | 2.51E-35 | Increased in Severe OA |
| | Binding of professional phagocytic cells | 8.25E-35 | Increased in Severe OA |
| | Interaction of phagocytes | 1.31E-34 | Increased in Severe OA |
| | Aggregation of cells | 2.08E-34 | Increased in Severe OA |
| | Endocytosis by eukaryotic cells | 2.86E-33 | Increased in Severe OA |
| | Cell spreading | 2.22E-32 | Increased in Severe OA |
| | Organisation of actin cytoskeleton | 1.34E-31 | Increased in Severe OA |
| | Cell movement of granulocytes | 2.09E-31 | Increased in Severe OA |
| | Cell movement of neutrophils | 3.42E-31 | Increased in Severe OA |
| | Aggregation of blood cells | 5.13E-31 | Increased in Severe OA |
| | Cell transformation | 6.51E-30 | Increased in Severe OA |
| | Migration of phagocytes | 2.55E-29 | Increased in Severe OA |
| | Phagocytosis | 3E-29 | Increased in Severe OA |
| | Cell movement | 3.26E-63 | Increased in mild OA |

| | | | |
|---|---|---|---|
| Mild OA vs healthy | Migration of cells | 4.29E-54 | Increased in mild OA |
| | Adhesion of immune cells | 3.01E-47 | Increased in mild OA |
| | Cell movement of blood cells | 1.98E-42 | Increased in mild OA |
| | Leukocyte migration | 3.23E-42 | Increased in mild OA |
| | Organisation of cytoskeleton | 1.04E-41 | Increased in mild OA |
| | Organisation of cytoplasm | 2.85E-41 | Increased in mild OA |
| | Cell movement of leukocytes | 1.22E-40 | Increased in mild OA |
| | Development of vasculature | 5.48E-39 | Increased in mild OA |
| | Cell movement of phagocytes | 4.4E-38 | Increased in mild OA |
| | Organismal death | 1.03E-37 | Decreased in mild OA |
| | Angiogenesis | 6.86E-37 | Increased in mild OA |
| | Cell movement of myeloid cells | 2.1E-36 | Increased in mild OA |
| | Vasculogenesis | 2.51E-35 | Increased in mild OA |
| | Binding of professional phagocytic cells | 8.25E-35 | Increased in mild OA |
| | Interaction of phagocytes | 1.31E-34 | Increased in mild OA |
| | Microtubule dynamics | 3.62E-33 | Increased in mild OA |
| | Cell movement of granulocytes | 2.09E-31 | Increased in mild OA |
| | Cell movement of neutrophils | 3.42E-31 | Increased in mild OA |
| | Cell transformation | 6.51E-30 | Increased in mild OA |
| | Invasion of cells | 1.17E-29 | Increased in mild OA |
| | Chemotaxis | 1.45E-29 | Increased in mild OA |
| | Adhesion of phagocytes | 1.09E-27 | Increased in mild OA |
| | Activation of cells | 9.89E-27 | Increased in mild OA |

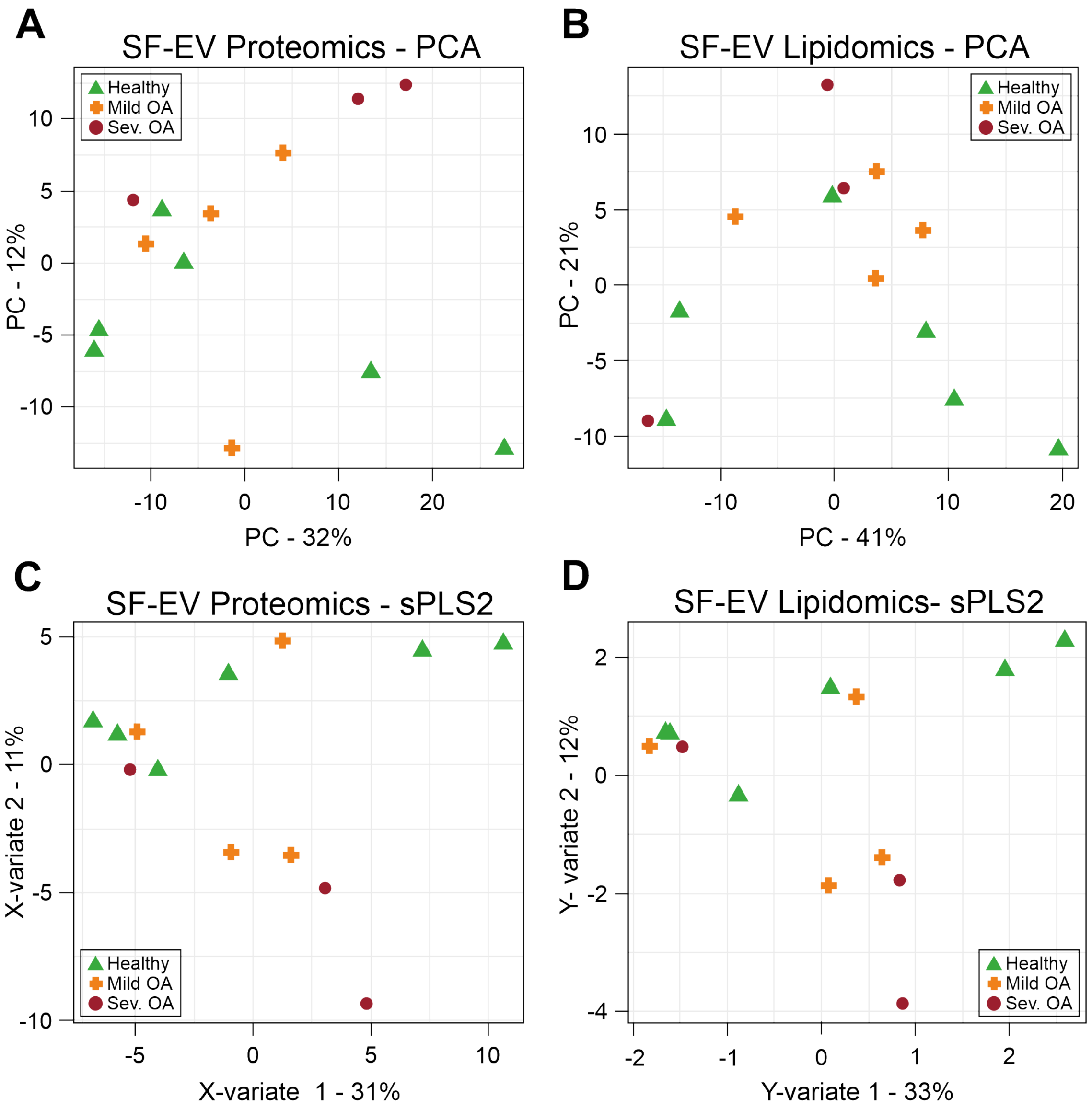


**Suppl. Fig. 3 : Normalisation of proteomics and lipidomics data for integrated analysis.** Healthy SF-EVs (green triangle, n=6), mild OA SF-EVs (orange cross, n=4), and severe OA SF-EVs (Sev. OA; red circle, n=3). Samples of both datasets were normalised by the sum. **A-B)** Principal component analysis (PCA) of the proteomics (A) and lipidomics (B) SF-EV datasets. **C-D)** Unsupervised multivariate Sparse Partial Least Squares regression (sPLS2) of the SF-EV samples in X-variate (for proteomics (C)) and Y-variate (for lipidomics (C)) components projected into the space spanned to the respective dataset.

**Table 5:** Candidate proteins for composite OA biomarker discovery.

| | **Protein** | **Membrane or cargo protein** | **Origin** | **Function** |
|---|---|---|---|---|
| | | **Group 1*– Downregulated proteins** | | |
| **Enzymes (oxidative stress)** | Peroxiredoxin 1 | Cargo [7] | Dendritic cells [8], spleen cells [9], osteoblasts [10], macrophages [11], thymus [12], pancreatic-$\beta$ cells [13], oligodendrocytes [14], | Modulation of redox signalling events for cellular protection [15] |
| **Membrane proteins** | CD163 (Scavenger receptor cysteine-rich type 1 protein M130) | Transmembrane [16] | Macrophage [16; 17] | Receptor for clearance of oxidative and proinflammatory haemoglobin/haptoglobin complexes, stimulator of heme-oxygenase-1 and anti-inflammatory heme metabolites [16] |
| **Membrane traffic** | Caveolin 1 | Cargo [18] | Ubiquitously expressed in all cell types [19] | Generation of caveolae (membrane curvatures) [18] |
| **Structural and cytoskeleton-related molecules** | Myosin regulatory light polypeptide 9 | Cargo [20] | Smooth muscle, spleen, fibroblasts (all non-muscle tissues) [20], Astrocytes [21], hippocampus [22] cancer [23; 24] | Modulation of contractile activity of smooth muscle and non-muscle cells by phosphorylation [25] |
| | Tubulin alpha chain | Cargo [26] | Ubiquitously expressed in all cell types [26] | Cellular morphology formation and maintenance, and intracellular transport, chromosomal segregation [26] |
| **Chaperones** | Heat shock protein 90 alpha | Cargo [27] | Fibroblasts [28], pancreatic-$\beta$ cells [13], oligodendrocytes [14], B cell [29], cancer [**?** ] | Protein folding, maturation, and involvement in signal transduction and transcriptional regulation [34] |
| **Endocytosis, exocytosis and trafficking** | Annexin A8 | Cargo [35; 36] | Chondrocytes [36], Leukocytes [37], osteoclasts [38], lung, liver, kidney, skin, placenta, and cornea [39] | Controls sorting and transport of late endosomes [35], chondrocyte differentiation marker under normal endochondral ossification [36] |
| **Signal transduction** | Protein S100-A4 | Cargo [40], secreted [41] | Chondrocytes [40], macrophages [41], fibroblasts, CD8+ T cells, monocytes, and eosinophils, cancer [42] | Regulation of calcium homeostasis, cell growth and motility, cell differentiation, and cell survival [40], promotion and induction of metastasis [42] |
| | | **Group 2** – Upregulated proteins** | | |

| | | | | |
|---|---|---|---|---|
| **Metabolism** | Aldo-keto reductase family 7 member A2 | Cargo | Cartilage [43], cardiomyocytes [44], brain, heart, kidney, liver, lung, prostate, skeletal muscle, small intestine, spleen [45] | Reduction of aldehydes and ketones to alcohols and metabolism of toxic aldehydes [44] |
| **Structural and cytoskeleton-related molecules** | Erythrocyte membrane protein band 4.1. like 2 | Cargo [46] | B cells [47], endothelial cells[48], Dendritic cells [49], Mesenchymal stem cells [50], T cells [51] | Role in the attachment of cytoplasmic proteins to the membrane, part of the FERM complex (composed of the 4.1 protein, ezrin, radixin, and moesin) [46] |
| | Ezrin | Cargo [52] | Chondrocytes [53], fibroblast [54; 55], B cells [47], milk [56], Dendritic cells [49], oligodendrocytes [14], cancer cells [57; 58], macrophages [59], monocytes [60], T cells [61], synovial fluid [62] B cell [47; 64], milk [56], | Cross-linker between the actin cytoskeleton and the plasma membrane, part of the ERM complex (ezrin/radixin/moesin), also involved in signal transduction cell migration, and survival [52] |
| | Moesin | Cargo [63] | Dendritic cells [49], endothelial cells [65], neutrophils [66], synovial fluid [62] | Links cytoplasmic regions of integrins and modulate their function [67; 72] |
| | Fermitin family member 3 | Cargo [67] | B cells [47], cancer cells [68; 69], Dendritic cells [49], endothelial cells [65], monocytes [60], neutrophils [66], red blood cells [70], platelets [71] | Bind to the cytoplasmic regions of integrins and modulate their function [67; 72] |
| | Actin alpha 2, smooth muscle (($\alpha$)-2 smooth muscle actin) | Cargo | Chondrocytes [73], monocytes and macrophages [74], fibroblasts [**?** ] | Actin isoform contributes to cell-generated mechanical tension, cell structure, tissue remodelling and contraction [77]. Role in fibrosis [**?** ] |
| | Vasodilator stimulated phosphoprotein | Cargo | B cells [47; 68], milk [56], dendritic cells [49], endothelial cells [65], monocytes [60], platelets [71; 83], T cells [61], cancer cells [69; 84; 85] | Associated with cell differentiation, mobility and tumour metastasis [82] |
| **Membrane proteins** | CD109 | Transmembrane [86] | Cancer cells [69; 84; 85; 87], T cells [61], Dendritic cells [49], Mesenchymal stem cells [50], endothelial cells [48], synovial tissue [88] | Modulation of pathological processes, such as osteoporosis, fibrosis and tumour metastasis [88], TGF-$\beta$ co-receptor and signalling inhibitor of TGF-$\beta$ in keratinocytes [89] |

| | | | | |
|---|---|---|---|---|
| | Thy-1 antigen/CD90 | Transmembrane [90] | T cells, NK cells, innate lymphoid cells [91], fibroblasts [92], Mesenchymal stem cells [50], cancer cells [69], endothelial cells, epithelial cells, neurons [90] | Involved in cancer development and metastasis, cell proliferation, differentiation, cell migration, apoptosis, mechanotransduction and cell adhesion[90; 93] |
| **Endocytosis, exocytosis and trafficking** | RAB8B, member RAS oncogene family | Cargo and secreted | Sperm[94], glioma cells [95], and SARS-CoV-2 sera [96] | Involved in membrane trafficking and the establishment of the Golgi apparatus [94] |
| **Signal transduction** | Rac family small GTPase 1 | Cargo and secreted | Hepatic cells [97], range of eukaryotic cells [98], equine synovial fluid and plasma [99] | Activated GTPases are involved in cellular proliferation, differentiation, motility, survival, and apoptosis [98] |
| **Transmembrane transport** | Potassium channel tetramerisation domain containing 12 | Membrane | Cancer cells[100; 101], Serum [102] | Involved in neuronal excitability through GABBA receptor signalling. Also found to suppress Wnt/Notch signalling, stem cell factors, and chromatin remodelers [103]. Associated with tetramerisation and gating of ion channels, cytoskeleton regulation, and transcriptional repression [104] |

* Proteins from group 1 correlated to the following phospholipids: PC 34:4, PI 38:6, PI 32:1.

** Proteins from group 2 correlated to the following phospholipids: PC O-32:3, SM 36:0;2, SM 41:3,2.
